# Systems Level Identification of a Matrisome-Associated Macrophage Polarization State in Multi-Organ Fibrosis

**DOI:** 10.1101/2022.12.20.521178

**Authors:** Kevin Y. Huang, Kunal Mishra, Harry Park, Xie Yi, John F. Ouyang, Enrico Petretto, Jacques Behmoaras

## Abstract

Tissue fibrosis affects multiple organs and involves a master-regulatory role of macrophages which respond to an initial inflammatory insult common in all forms of fibrosis. The recently unraveled multi-organ heterogeneity of macrophages in healthy and fibrotic human disease suggest that tissue resident macrophages, expressing osteopontin (SPP1), associate with lung and liver fibrosis. However, the conservation of this SPP1+ macrophage population across different tissues, and its specificity to fibrotic diseases with different etiologies remain unclear. Integrating 13 single cell RNA-sequencing datasets to profile 225,985 tissue macrophages from healthy and fibrotic heart, lung, liver, kidney, skin and endometrium, we extended the association of SPP1+ macrophages with fibrosis to all these tissues. We also identified a subpopulation expressing matrisome-associated genes (e.g., matrix metalloproteinases and their tissue inhibitors), functionally enriched for ECM remodeling and cell metabolism, representative of a matrisome-associated macrophage (MAM) polarization state within SPP1+ macrophages. Importantly, the MAM polarization state follows a differentiation trajectory from SPP1+ macrophages, which was conserved across all fibrotic tissues and driven by NFATC1 and HIVEP3 regulons. Unlike SPP1+ macrophages, the MAM polarization state shows a positive association with ageing in mice and humans, and across multiple tissues during homeostasis. These results suggest an advanced, age-dependent polarization state of SPP1+ macrophages in fibrotic tissues as a result of prolonged inflammatory cues within each tissue microenvironment.

## INTRODUCTION

Fibrosing diseases comprise a multitude of human organs and share a common end-point: unresolved inflammation characterized by abnormal production of extracellular matrix (ECM) and interstitial scar formation. In almost all tissues, macrophages control the pathobiology of fibrosis in a timely manner. Macrophage polarization accompanies the progressively changing tissue microenvironment and macrophages modulate fibroblast activation and ECM-producing myofibroblast *trans*-differentiation through soluble factors or direct cell-cell interaction [1–4]. The outcome of fibrosis is directly dependent on tissue macrophage heterogeneity as different macrophage subpopulations can have contrasting modulatory effects on fibrogenesis [5]. For instance, in the lung and heart, monocyte-derived macrophages infiltrate the inflamed tissues and show pro-fibrogenic activity [6, 7]. On the contrary, tissue resident macrophages negatively regulate fibrosis as their loss exacerbates cardiac and lung fibrosis [8, 9].

Although fibrotic lesions may occur in distinct anatomical sites within the same organ, the process of fibrogenesis is a shared hallmark of several diseases, including heart failure, chronic kidney disease, liver cirrhosis and interstitial lung disease (ILD) [10]. In addition, fibrosis of the skin and uterus are often seen in patients diagnosed with systemic sclerosis (SSc) [11] and endometriosis [12, 13], respectively. All of these conditions also share an inflammatory component with macrophage involvement in affected organs. Thus, knowledge of homeostatic (disease-free) tissue macrophage heterogeneity is required to understand macrophage subpopulations and activation states during fibrosis in these etiologically different diseases. To this aim, cross-tissue human single cell atlas initiatives provided a useful resource to address the complexity of homeostatic tissue resident macrophage states at an unprecedented resolution in multiple organs [14–16]. In addition to these resources, recent single cell transcriptomics studies compared healthy and fibrotic (or inflammatory) human tissues, and further revealed the evolution of the macrophage activation states during inflammation/fibrosis in multiple organs [17–28].

Three main findings emerged from these large-scale, single cell-based resources for healthy and fibrotic human tissues: (i) the heterogeneity of macrophages is conserved across human tissues with tissue-restricted functionalization (e.g. expression of genes related to iron recycling in erythrophagocytic macrophages of spleen and liver Kupffer cells [14]); (ii) a core tissue-resident macrophage transcriptome broadly associates with lipid-related pathways during homeostasis in multiple human tissues [15]; (iii) among other cell markers, a core macrophage population express the matricellular glycoprotein osteopontin (*SPP1*), a gene implicated in the development of wound healing and fibrosis [29–32]. Importantly, SPP1+ macrophages (also termed as scar-associated macrophages – SAMs, initially described in hepatic fibrosis [33] and further refined at a single cell level in cirrhotic liver [25]) have been described as pro-fibrotic cells in human pulmonary and hepatic fibrosis [24, 25, 27, 34]. However, the broader implication of SPP1+ SAMs in multi-organ fibrosis and their potential heterogeneity remain to be identified. Furthermore, SPP1+ SAMs share an overlapping transcriptome with TREM2+ tumor-associated macrophages (TAMs) [35], lipid-associated macrophages (LAMs) [36], and disease-associated microglia (DAM) [37, 38], – hence lack specific functionality for tissue fibrosis.

Here we focused on human tissues and analyzed scRNA-seq data from 13 studies carried out in healthy and disease tissues characterised by fibrosis of the heart, lung, liver, kidney, skin and uterus. Focusing on the tissue macrophage compartment (225,985 cells), we show that SPP1+ macrophages are indeed a prominent feature conserved in multi-organ fibrosis in humans. Stemming from SPP1 macrophages, we identified a functionally defined polarization state, called matrisome-associated macrophage (MAM), which we investigated in the context of fibrosis (in disease) and aging (in homeostasis).

## RESULTS

### SPP1+ macrophages increase during fibrotic disease across tissues in humans

Based on the previously established association of pro-fibrotic SPP1+ macrophages with cirrhotic liver and lungs from idiopathic pulmonary fibrosis (IPF) patients [24, 25], we hypothesized that this macrophage population can be detected in other human tissues and can associate with broader fibrotic disease state. Thus, we interrogated human single cell datasets which contain, among other cells, macrophages in healthy and fibrotic tissues. For consistency and ease of normalization for further meta-analysis, we focused on datasets generated in a single platform (10X), using live cell isolation protocols (Table 1). This led to a total of 225,985 tissue macrophages from healthy and diseased human liver, lung, heart, skin, endometrium, and kidney (Table 1). The pathologies affecting these organs included cirrhosis, IPF, other ILD, SSc, dilated cardiomyopathy (DCM), ischemic cardiomyopathy (ICM), keloid, endometriosis, chronic kidney disease (CKD) and acute kidney injury (AKI). The percentage of macrophages vary across the different tissues, ranging from 49% in the lung [17] to only 0.4% of total cells in the skin [39].

**Table 1.**
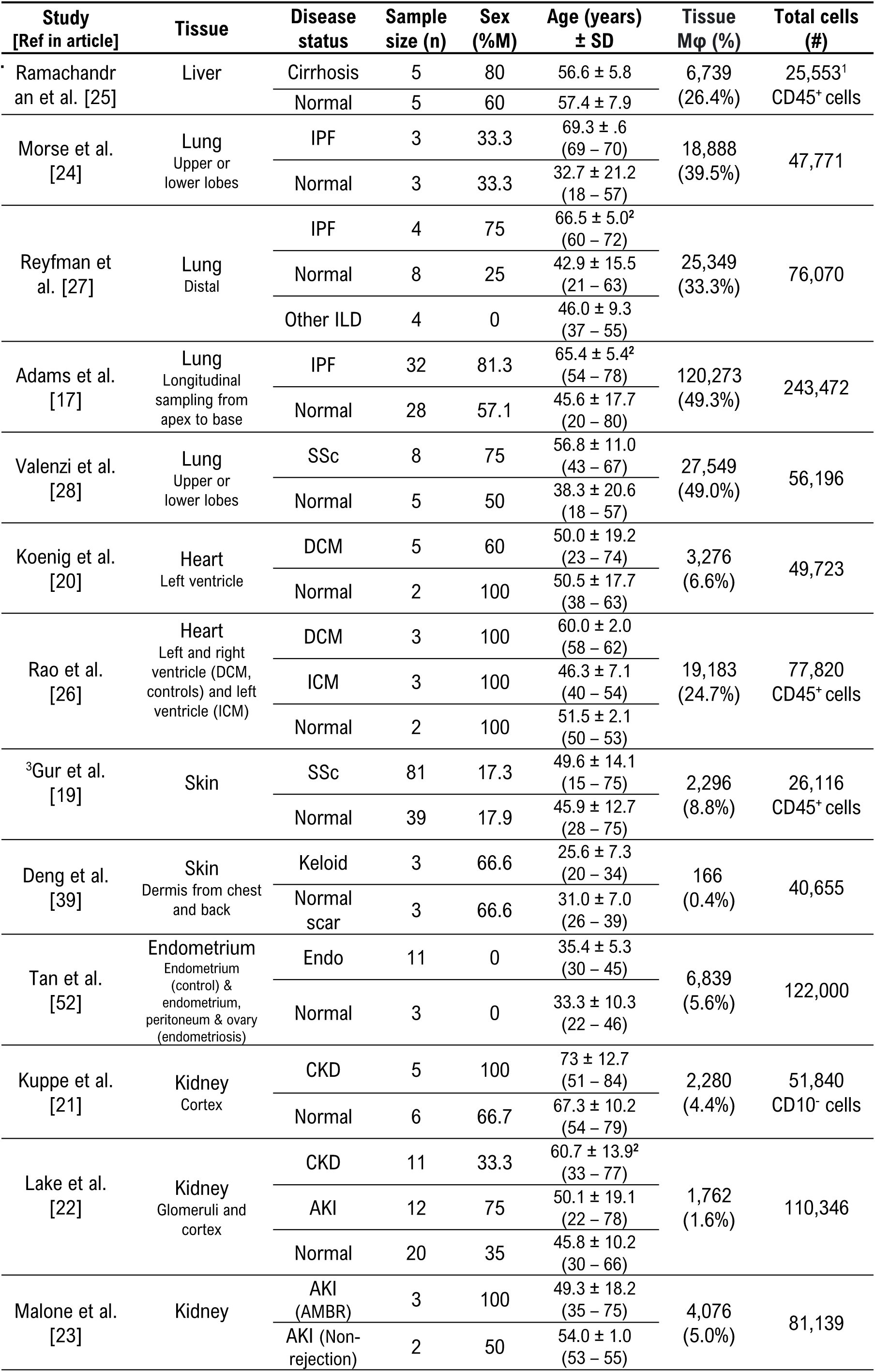
Summary of scRNA-seq datasets analyzed in this study. For each study, the sample size (number of single cell datasets), age/gender, disease status, the number (and percentage) of tissue resident macrophages (Mφ), and the total number of live cells sequenced are indicated. For studies that opted for positive or negative cell sorting methods (e.g., CD45+ cells), the total number of live cells is indicated with the cell isolation method. IPF: idiopathic pulmonary fibrosis, ILD: interstitial lung disease, SSc: systemic sclerosis; DCM: dilated cardiomyopathy; ICM: ischemic cardiomyopathy; Endo: endometriosis; CKD: chronic kidney disease; AKI: acute kidney injury; AMBR: antibody-mediated rejection. ^1^ The number of CD45+ cells has been derived from the dataset downloaded from GEO (otherwise this number has been provided in the original publication and reported here). ^2^ P<0.05 for the comparison of age between disease and control groups (two-tailed Mann-Whitney U test). ^3^ Massive parallel single-cell RNA-sequencing (Mars-seq) was used to profile single-cell transcriptomics instead of 10X Chromium kit.

We then performed data-driven clustering of macrophages within each tissue (see Methods Supplementary Figures 1-6, Supplementary Table 1). We obtained between 4-6 macrophage clusters on average across the six human tissues, with presence of SPP1+ macrophages as a common cell cluster throughout (Figure 1A-F). Compared with all macrophages, the SPP1+ macrophages are positively enriched for several processes, including ECM degradation/remodeling and metabolic processes such as oxidative phosphorylation (Supplementary Table 3). Compared with healthy control tissue, the proportions of SPP1+ macrophages were increased in IPF and SSc lungs (Figure 1A), cirrhotic liver (Figure 1B), DCM/ICM heart (Figure 1C), SSc and keloid skin (Figure 1D), endometriosis uterus (Figure 1E), and CKD/AKI kidney (Figure 1F). This was confirmed when the proportion of SPP1+ macrophages was assessed in individual patients, which showed an association with disease state in multiple conditions (Figure 1G), although endometriosis did not reach statistical significance. Amongst the top differentially expressed genes in SPP1+ macrophages, *SPP1* itself was found as the most and more consistently upregulated gene in disease across all 6 tissues (Figure 1H). These results confirmed and extended the association of SPP1+ macrophages with multi-organ fibrosis, and prioritized *SPP1* as the most suitable marker transcript of this macrophage subset.

**Figure 1.**
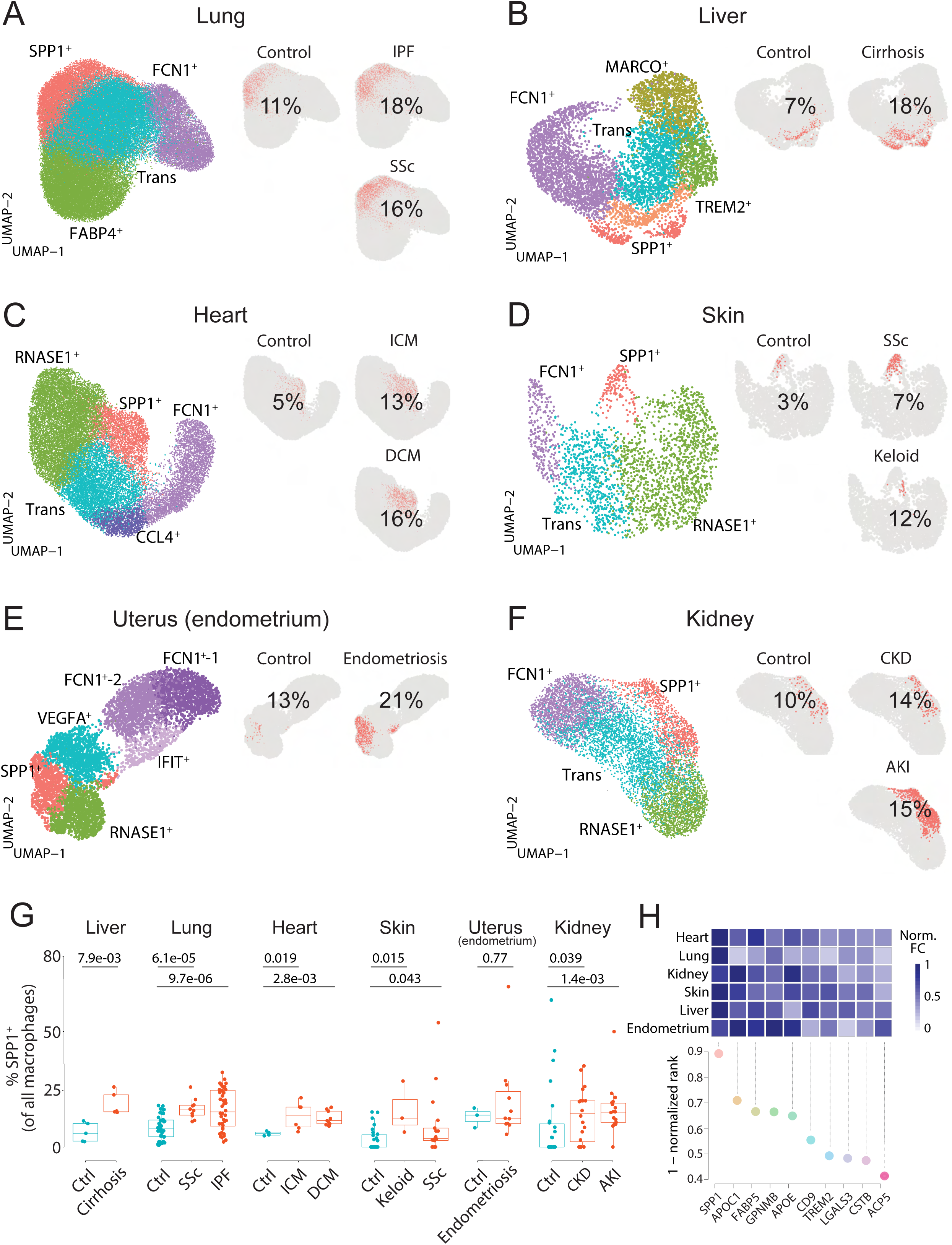
Resident SPP1+ macrophages are increased in fibrotic disease across-tissues. (**A-F**) Uniform Manifold Approximation and Projection (UMAP) dimensionality reduction of total macrophages in scRNA-seq of the lungs (**A**), liver (**B**), heart (**C**), skin (**D**), endometrium (**E**), and kidney **(A)** (**F**) from fibrotic disease patients or controls (see Table 1 for details on samples and number of macrophages in each tissue). Clusters were denoted as SPP1+ macrophages, FCN1^+^ monocytes, homeostatic macrophages (RNASE1^+^, or MARCO+ macrophages), or transitional macrophage (“Trans” or “VEGF+” macrophages) based on expression of marker genes, see Supplementary Table 1 and Supplementary Figure 1-6 for details. SPP1+ macrophages in each tissue are stratified by disease conditions and coloured in orange on separate UMAP; the proportion of SPP1+ macrophages out of all macrophages in each condition is indicated. (**G**) Boxplot summarizing the relative proportion of SPP1+ macrophages (out of all macrophages) in each subject stratified by disease status. Liver: (control) n = 5, (cirrhosis) n = 5; lung: (control) n = 44, (IPF) n = 44, (SSc) n = 10; heart: (control) n = 4, (ICM) n = 6, (DCM) n = 9; skin: (control) n = 27, (keloid) n = 3, (SSc) n = 17; uterus (endometrium): (control) n = 3, (endometriosis) n = 11; kidney: (control) n = 26, (CKD) n = 20, (AKI) n = 18. Colour of dots delineates cases (orange) and control (blue). The Wilcoxon rank-sum test was used to evaluate the significance of the difference between groups. (**H**) Heatmap displaying the fold change (FC) in expression of the top differentially expressed genes (DEGs) (x-axis) upregulated in SPP1+ macrophage compared to other macrophages in different tissues (top panel). For each gene, the FC in expression was scaled to the highest fold change in each tissue. Rank-plot prioritizing the top DEGs that are conserved across tissues, where the *x-axis* are the marker genes and *y-axis* is the cumulative rank of each DEG based on FC within each tissue (bottom panel).

### A matrisome-associated macrophage state of polarization within SPP1+ cells

We next reasoned that SPP1+ macrophages may be heterogeneous and contain polarization states that can be used for a refined functional characterization of these cells in multi-organ fibrosis. Single-cell transcriptomics capture a continuum of macrophage phenotypes within patient tissues who often show differences in disease severity. As such, the transcriptional resolution of a cell state in a single tissue can be confounded by several factors such as sample size, sequencing depth, disease heterogeneity, etc. We thus performed an integrative meta-analysis of SPP1+ macrophages throughout 6 tissues including 9 fibrotic conditions and their matched controls. Using unsupervised clustering methods in SPP1+ macrophages (Supplementary Figure 7a-c) we identified three subpopulations, including a matrisome-associated macrophage (MAM) subcluster (defined as SPP1+MAM+) and other two subclusters enriched for inflammatory and ribosomal processes (Figure 2A and Supplementary Figure 7d,e). The relative distribution of macrophages in the tissues of origin was not associated with the subcluster identification within the SPP1 macrophages (Supplementary Figure 7f). For clarity of nomenclature, we refer to SPP1+ macrophages as those defined in multiple tissues during multi-organ fibrosis (Figure 1 and Supplementary Figures 1-6), to SPP1+MAM+ as the cells derived from re-clustering of SPP1+ macrophages in the single cell meta-analysis (Figure 2A and Supplementary Figure 7) and to SPP1+MAM-as the population of SPP1+ macrophages devoid of SPP1+MAM+, i.e., clusters 0 and 2 within SPP1+.

**Figure 2.**
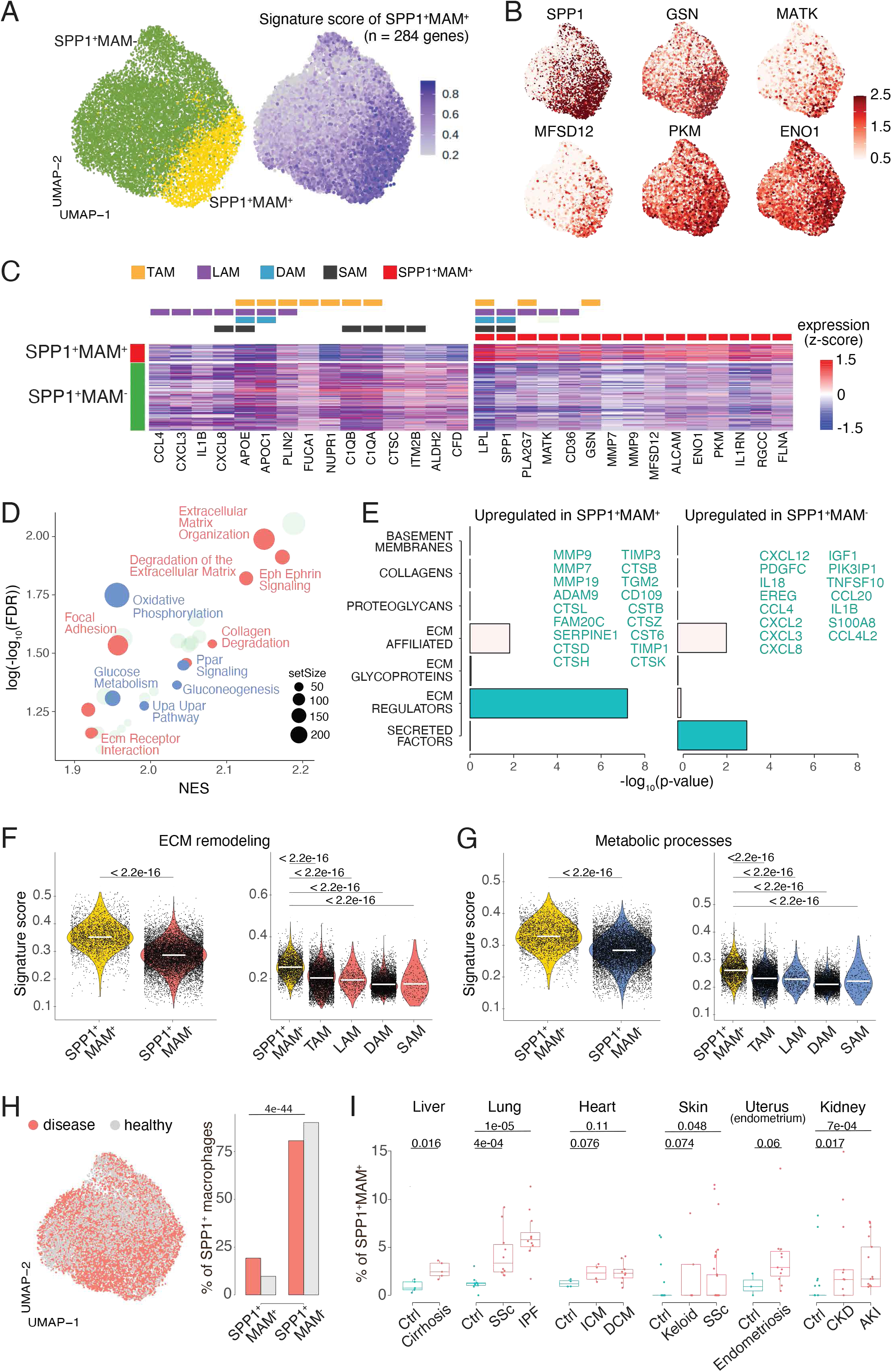
Identification of a matrisome-associated macrophage (MAM) state within SPP1+ macrophages. (**AA)** UMAP dimensionality reduction of all SPP1+ macrophages merged from the lung, liver, heart, skin, kidney, and endometrium tissues following Seurat scRNA-seq datasets integration (see Methods). Unsupervised clustering of SPP1+ macrophages identified a matrisome-associated macrophage (MAM) subpopulation (yellow), defined as SPP1+MAM+ (left panel; Supplementary Figure 8). Each macrophage is coloured by the expression level of SPP1+MAM+ signature genes (n=284 genes, reported in Supplementary Table 4; right panel). (**B**) UMAP of SPP1+ macrophages with each cell coloured by the expression of top 6 markers for SPP1+MAM+ as predicted by COMET (see Methods and Supplementary Figure 9). (**C**) Heatmap showing the scaled gene expression levels of marker genes for SPP1+MAM+ and SPP1+MAM-macrophages; marker genes were selected based on highest log2(fold change) between SPP1+MAM+ and SPP1+MAM-cells. Markers of SPP1+MAM+ (shown in short as MAM), tumor-associated macrophages (TAMs) [35], lipid-associated macrophages (LAMs) [15, 36], disease-associated microglia (DAM) [37, 38], and scar-associated macrophages (SAMs) [25, 34] are indicated above the heatmap. (**D**) Scatterplot summarizing the results of GSEA analysis of differentially expressed genes (DEG) between SPP1+MAM+ and SPP1+MAM-macrophages. For each significant pathway (FDR<5%), *x-axis* indicates the normalized enrichment score (NES) and *y-axis* the significance of the enrichment; the size of each dot is proportional to the number of leading-edge genes in the enrichment. Metabolism-related pathways are coloured in blue, ECM remodelling in red, and homeostatic/other terms in green. (**E**) Barplots summarizing pathway enrichment of DEG upregulated in SPP1+MAM+ (left panel) or in SPP1+MAM-macrophages (right panel) using a matrisome-specific database [41]. Filled bars indicate significant enrichment (P<0.05), determined by hypergeometric test. Genes up-regulated in SPP1+MAM+ or SPP1+MAM-macrophages and the corresponding enriched pathway are displayed. (**F-G**) Violin plots showing the transcriptomic signature scores (*y-axis*) of SPP1+MAM+ ECM remodelling gene set (n=42 genes, Supplementary Table 6) (**F**), and SPP1+MAM+ metabolic processes gene set (n=53 genes, Supplementary Table 7) (**G**) in SPP1+MAM+ and SPP1+MAM-macrophages (*left*), and in other tissue resident macrophages with overlapping transcriptomic signatures (*right*). SPP1+MAM+, n = 2,729; TAMs, n = 6,556; LAMs, n = 602; DAM, n = 5,645; SPP1+MAM-, n = 10,971; SAMs, n = 442 cells. (**H**) UMAP dimensionality reduction of SPP1+ macrophages from all tissues, where each cell is coloured according to the disease status (*left*). Proportion of SPP1+MAM+ or SPP1+MAM-out of all SPP1+ macrophages from disease or healthy control samples. The overrepresentation of SPP1+MAM+ cells in disease was evaluated by hypergeometric test and indicated. (**I**) Boxplot showing the percentage of SPP1+MAM+ in all macrophages, stratified by tissue and patient status (orange, disease; blue, control). Liver: (control) n = 5, (cirrhosis) n = 5; lung: (control) n = 13, (SSc) n = 10, (IPF) n = 10; heart: (control) n = 4, (ICM) n = 6, (DCM) n = 9; skin: (control) n = 27, (keloid) n = 3, (SSc) n = 17; uterus (endometrium): (control) n = 3, (endometriosis) n = 11; kidney: (control) n = 18, (CKD) n = 20, (AKI) n = 18. Unless otherwise indicated, a two-sided Wilcoxon rank-sum test was used to evaluate the significance of differences between groups.

We defined a SPP1+MAM+ transcriptomic signature comprising 284 genes (Supplementary Table 4). In order to identify markers that delineate SPP1+MAM+ polarization state in an unbiased way, we used COMET [40] (Supplementary Figure 8 and see methods) and found *GSN*, *MATK*, *MFSD12*, *PKM* and *ENO1*, in addition to *SPP1* (Figure 2B). Because of the transcriptional overlap between the previously defined macrophage states such as LAMs [36], TAMs [35] DAM [37, 38] and SAMs [25], we next performed systematic comparative analyses between SPP1+MAM+ and SPP1+MAM- (our meta-analysis) in relation to the previously described macrophage activation states (Figure 2C, and See Supplementary Figure 9 for derivation of DAM signature). This analysis revealed *MMP7*, *MMP9*, *ALCAM*, *ENO1*, *RGCC,* and *FLNA* as genes uniquely differentially expressed and upregulated in association with the SPP1+MAM+ polarization state (Figure 2C). As expected, some markers (e.g., *LPL*, *SPP1*, *APOE*, *APOC1*) were shared across LAMs, DAM and TAMs, and as such, did not add context specificity to the SPP1+MAM+ (Figure 2C).

To achieve further transcriptional specificity, we reasoned that the activation of matrisome-related genes polarizes the SPP1+ macrophages further and that without the MAM signature, the cells will recap the transcriptional state of the previously identified SAMs [25] (Supplementary Figure 10a,b). We indeed found a positive correlation between pseudo-bulk profiles of SAMs (as defined by [25]) and SPP1+MAM-macrophages, while SAMs were negatively correlated (i.e., were dissimilar) with SPP1+MAM+ macrophages (Supplementary Figure 10c). In keeping with this, the markers of SAMs were not differentially expressed in the SPP1+MAM+ macrophages (Supplementary Figure 10d). This suggested that SPP1+MAM+ are enriched for newly acquired pathways that may be relevant for fibrosis. When compared to SPP1+MAM-, gene set enrichment analysis (GSEA) of differentially expressed genes showed predominance of ECM remodeling and cell metabolism related pathways in SPP1+MAM+ (Figure 2D, Supplementary Table 5). Interestingly, when we queried a proteomics-based matrisome database [41], pathway enrichment for SPP1+MAM+ and SPP1+MAM-genes was distinctive, with SPP1+MAM+ showing an association with ECM regulators and SPP1+MAM-with secreted factors (Figure 2E). Specifically, SPP1+MAM+ are characterized by expression of MMPs (*MMP7*, *MMP9*, *MMP19*), TIMPs (*TIMP1*, *TIMP3*), cathepsin family genes with endopeptidase activity (*CTSB*, *CTSL*, *CTSH*, *CTSD*, *CTSZ*) – suggesting these cells represent an overall macrophage state with active ECM remodelling. On the other hand, SPP1+MAM-are enriched specifically for secreted factors and cytokine-cytokine receptor signaling (Figure 2E), as well as for interferon/complement and immune related processes (Supplementary Table 5). This suggests a phenotypically distinct state of activation of SPP1+MAM+. Since the “ECM remodeling” and “metabolic processes” define predominantly SPP1+MAM+, we compared the presence of these two signatures (Supplementary Tables 6-7) throughout the previously reported macrophage activation states (Figure 2F and G). SPP1+MAM+ showed significant up-regulation of ECM remodeling and metabolic processes gene signatures compared to SPP1+ macrophages (Figure 2F and G). When compared to TAMs, LAMs, DAM and SAMs, SPP1+MAM+ also showed an increased activation of the two processes (Figure 2F and G), confirming their distinct transcriptomic profile that may implicate this polarization state more specifically in ECM remodeling during multi-organ fibrosis. The SPP1+MAM+ cells were present in healthy and disease tissues, and the percentage of SPP1+MAM+ cells across all macrophages was doubled in disease compared with control cells (Figure 2H). Despite showing different statistical significance levels, SPP1+MAM+ cells were increased in patients with cirrhosis, SSc, IPF, ICM, DCM, keloid, CKD and AKI (Figure 2H). Unlike SPP1+ macrophages (P=0.77)(Figure 1G), SPP1+MAM+ showed borderline significance (P=0.074) for increased percentage in endometriosis compared with controls (Figure 2H).

### SPP1+MAM-macrophages polarize towards SPP1+MAM+ state in fibrotic disease

Having established the distinct transcriptional and functional signatures of SPP1+MAM+, we next investigated their relationship with the other macrophage populations and their differentiation state in the trajectory of these cells, especially with regard to SPP1+MAM-. Interestingly, slingshot differentiation trajectory analyses identified 2 main trajectories: one trajectory from the FCN1^+^ infiltrating monocytes/macrophages toward the homeostatic (RNASE1^+^ or MARCO+) and another trajectory from FCN1^+^ monocytes/macrophages toward SPP1+MAM+ (Figure 3A-F). The differentiation trajectories inferred by Slingshot were confirmed by Monocle (Supplementary Figure 11). In the liver, there were two homeostatic trajectories toward either RNASE1^+^ or MARCO+ (Figure 3B). Notably, in all the trajectories throughout the tissues and fibrotic diseases, SPP1+MAM+ were at the end of the trajectory preceded by SPP1+MAM-state (Figure 3A-F), confirming SPP1+MAM+ represent a common polarization state of SPP1+ macrophages. When stratified according to the disease status, the cell propensity score (CPS, i.e., the probability of SPP1+MAM-macrophages to differentiate into SPP1+MAM+) was significantly more prominent in the fibrotic disease state throughout all tissues (Figure 3A-G). In other words, compared with the SPP1+MAM-macrophages in controls, we found significantly more SPP1+MAM-macrophages with high propensity to differentiate into SPP1+MAM+ in each disease patient group. Interestingly, some healthy tissues, such as the heart, had a relatively elevated CPS compared with healthy liver, skin and endometrium, which showed negligible propensity of differentiation of SPP1+MAM-into SPP1+MAM+. This suggests that some of the “control” subjects might include tissue samples surgically removed from individuals with heart conditions such as arrhythmia or non-ischemic cardiomyopathy. Taken together, these results show an exacerbated SPP1+MAM+ differentiation in fibrotic disease tissues.

**Figure 3.**
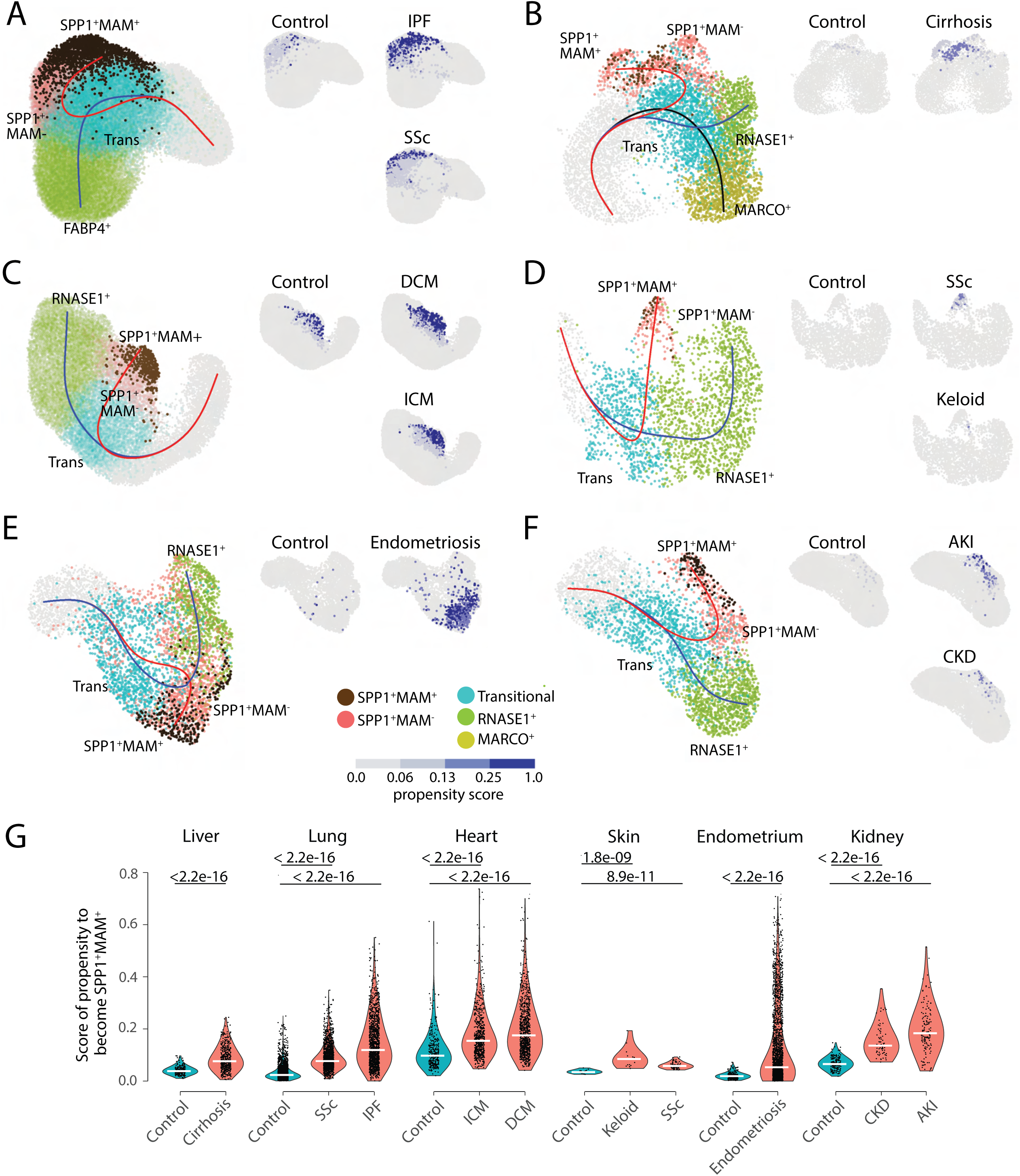
The differentiation trajectory from FCN1^+^ monocytes to SPP1+MAM+. (**A-F**) Slingshot differentiation trajectory analyses of macrophages in the lung (**A**), liver (**B**), heart (**C**), skin (**D**), endometrium (**E**), and kidney (**F**). The predicted trajectories are drawn on UMAP projection of macrophage, where the trajectory from FCN1^+^ monocyte to SPP1+MAM+ is indicated by a red line, and the one from FCN1^+^ monocytes to homeostatic macrophages (RNASE1^+^ or MARCO+) is indicated by a blue line. Macrophages are further stratified by disease and controls in each tissue and plotted separately on different UMAP, with each SPP1+MAM-macrophages coloured by its propensity to differentiate towards the SPP1+MAM+ state. To improve the visualization of the propensity score of each cell across all tissues, we binned the score into 4 bins with different shades of blue. (**G**) Violin plots displaying the SPP1+MAM-macrophages propensity score towards the SPP1+MAM+ state in each tissue, separated by disease conditions. Number of cells is as follows, liver: (control) n = 213, (cirrhosis) n = 471; lung: (control) n = 2,602, (SSc) n = 1,772, (IPF) n = 1,384; heart: (control) n = 327, (ICM) n = 800, (DCM) n = 900; skin: (control) n = 23, (keloid) n = 20, (SSc) n = 110; endometrium: (control) n = 69, (endometriosis) n = 1,090; kidney: (control) n = 608, (CKD) n = 80, (AKI) n = 658. Colour of violin delineates disease (orange) and control (blue). Unless otherwise indicated, a two-sided Wilcoxon rank-sum test was used to evaluate the statistical significance (P-value) of differences between two groups.

### Regulons driving the differentiation trajectory and activation of SPP1+MAM+ polarization state

Since SPP1+MAM+ are at the end-trajectory of different macrophage states and are likely to be a hallmark of multi-organ fibrosis, we next investigated transcription factor networks (regulons) that might be involved in their differentiation and activation. The SPP1+MAM+ state of polarization was associated with the likelihood of activation of several regulons (Figure 4A). Amongst these, NFATC1, HIVEP3, THRA, NR1H3, ETV5 and MAF regulons show the highest specificity for the signature of SPP1+MAM+ (Figure 4B). We then investigated the association between SPP1+MAM+ and the activity of each regulon during the differentiation trajectories from transitional (trans), SPP1+MAM- and SPP1+MAM+ states. To this aim, we fit a linear regression model to assess the SPP1+MAM+/regulon association along the two differentiation trajectories: (1) from trans toward SPP1+MAM- and (2) from SPP1+MAM- to SPP1+MAM+ (Figure 4C). Interestingly, this analysis showed that NFATC1 and HIVEP3 regulons were positively associated (Spearman *ρ* = 0.67-0.69) in the trajectory from SPP1+MAM- to SPP1+MAM+, and to a lower extent in the trajectory from trans to SPP1+MAM- (Spearman *ρ* = 0.25-0.35). In addition, the change in the regression slopes between the two differentiation trajectories (slope 0.48-0.49 *vs* 0.09- 0.19, Figure 4C) suggests a late activation of NFATC1 and HIVEP3 regulons throughout the differentiation trajectory, concordant with the emergence of the MAM+ polarization state. On the contrary, THRA, NR1H3, ETV5 and MAF were regulons that were associated from transition toward the SPP1+MAM- as well as from SPP1+MAM- to SPP1+MAM+ states, and with similar regression slopes (Figure 4C). The activities of NFATC1 and HIVEP3 regulons were positively associated with the SPP1+MAM+ polarization state when all SPP1+ macrophages were considered (Figure 4D). Furthermore, the regulon activity in SPP1+MAM+ was increased with respect to the activity in homeostatic macrophages (Figure 4E). Taken together, these results prioritize NFATC1 and HIVEP3 regulons as driving the differentiation of SPP1+MAM- macrophages towards the SPP1+MAM+ polarization state.

**Figure 4.**
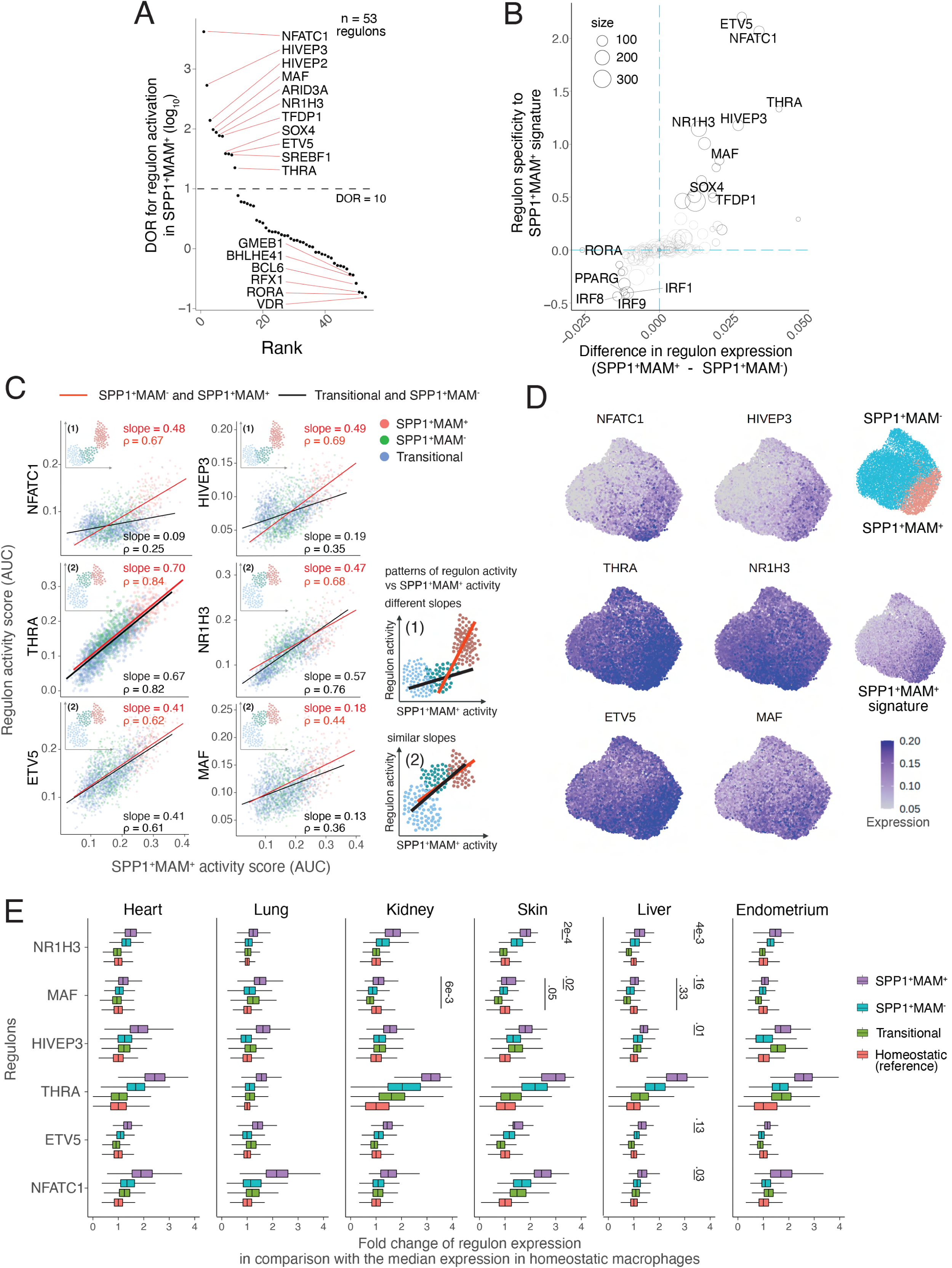
A core set of regulons drives the polarization of SPP1+MAM- to SPP1+MAM+. (**AA)** Rank plot for the top 53 regulons active in SPP1+MAM+ macrophages and driving the differentiation from SPP1+MAM- to SPP1+MAM+ macrophages. For each regulon, the diagnostic Odd Ratio (DOR) (*y- axis*) calculates the odds of being activated in a SPP1+MAM+ over the odds of being activated in a SPP1+MAM- macrophages (see Methods for details). A positive DOR is associated with the regulon activation in SPP1+MAM+, and a DOR = 10 is usually associated with good positive test (dashed line). DOR values are reported as log10(DOR). (**B**) Scatterplot plots the specificity of regulon genes to the transcriptomic signature of SPP1+MAM+ (*y-axis*) against the difference in expression of a regulon between SPP1+MAM+ and SPP1+MAM- macrophages (*x-axis*). Briefly, the specificity score of the regulon reflects the degree of overlap between regulon genes and the SPP1+MAM+ signature genes (see **Methods** for details). (**C**) Scatterplot showing each cell’s activity score of selected regulons (*y-axis*) against the cell’s SPP1+ MAM+ transcriptomic signature score (*x-axis*). These scores are shown for all the transitional, SPP1+MAM- and SPP1+ MAM+ macrophages pulled together across tissues, and each dot on the scatterplot is a macrophage coloured by its cell-type identity. For each regulon, separate linear regression models have been fitted to each trajectory: (*i*) from the transitional to SPP1+MAM- macrophages (black regression line), and (*ii*) from SPP1+MAM- to SPP1+MAM+ macrophages (red regression line). We defined two patterns of change in the regulon activity score: pattern (1) where there is a change in the regression slopes between the two differentiation trajectories, and pattern (2) with no apparent change in the regression slopes (see illustrations on the right). For each regulon, the pattern is indicated in the scatterplot (insert), and the slope of the linear regression and the Spearman correlation (ρ) are indicated. (**D**) UMAP of SPP1+ macrophages coloured by expression level of selected regulons, and by the expression of the SPP1+MAM+ signature. (**E**) Box plots summarizing the fold change of regulon gene expression from SPP1+MAM+, SPP1+MAM-, transitional macrophages compared with regulon expression in homeostatic macrophages. Within each tissue, the ratio (fold change, FC) between the regulon expression (i.e., AUC score by pySCNEIC, see Methods) in each cell from SPP1+MAM+, SPP1+MAM-, or transitional macrophage populations with respect to the median expression in the homeostatic macrophages was calculated. The box plots show the median, interquartile range and lowest and highest values (whiskers) of the FCs across all cells (*x-axis*). The statistical significance (P- value) of the differences in FCs in regulon expression between SPP1+MAM+ macrophages and other macrophages (i.e., SPP1+MAM+ *vs* SPP1+MAM-, SPP1+MAM+ *vs* transitional Mac, or SPP1+MAM+ *vs* homeostatic macrophages) were evaluated using a two-tailed Wilcoxon rank-sum test. Unless otherwise indicated in the figure, the P-value < 0.0001 for each comparison of regulon expression with SPP1+MAM+ macrophages.

### SPP1+MAM+ polarization state is associated with aging in healthy tissues

Physiological aging is one of the risk factors for fibrotic disease [42, 43] which is often associated with impaired resolution of the prior inflammatory insults [44] and metabolically activated macrophages [45, 46]. We next considered the tissue macrophages from the homeostatic (disease-free) state, and evaluated the SPP1+MAM+ signature activity as a function of age in heart and lung tissues of control individuals used in comparison with fibrotic disease tissues (Figure 1). We found a significant positive correlation between SPP1+MAM+ activity in healthy lung and heart and age (*P*=0.002), which was not the case with SPP1+MAM- state (Figure 5A and B, and Supplementary Figure 12, Supplementary Figure 13A). We extended this analysis by adding healthy skin macrophages (Supplementary Figure 13B). Despite the significantly smaller number of cells in skin (cells across individuals: 5-57 in skin *vs* 202- 5928 in lung and 311-1248 in heart), and which resulted in a significant effect of tissue-variability for skin, we still found a significant association between SPP1+MAM+ activity and age (*P*=0.005) when we combined all three tissues in the analysis (Supplementary Figure 13C). To strengthen and extend this association to more tissues, we used a single cell transcriptomic atlas of young and old mice [47], which confirmed increased expression of SPP1+MAM+, SPP1+MAM+ metabolic and SPP1+MAM+ ECM signatures in macrophages from kidney and lungs in old mice (Figure 5C).

**Figure 5.**
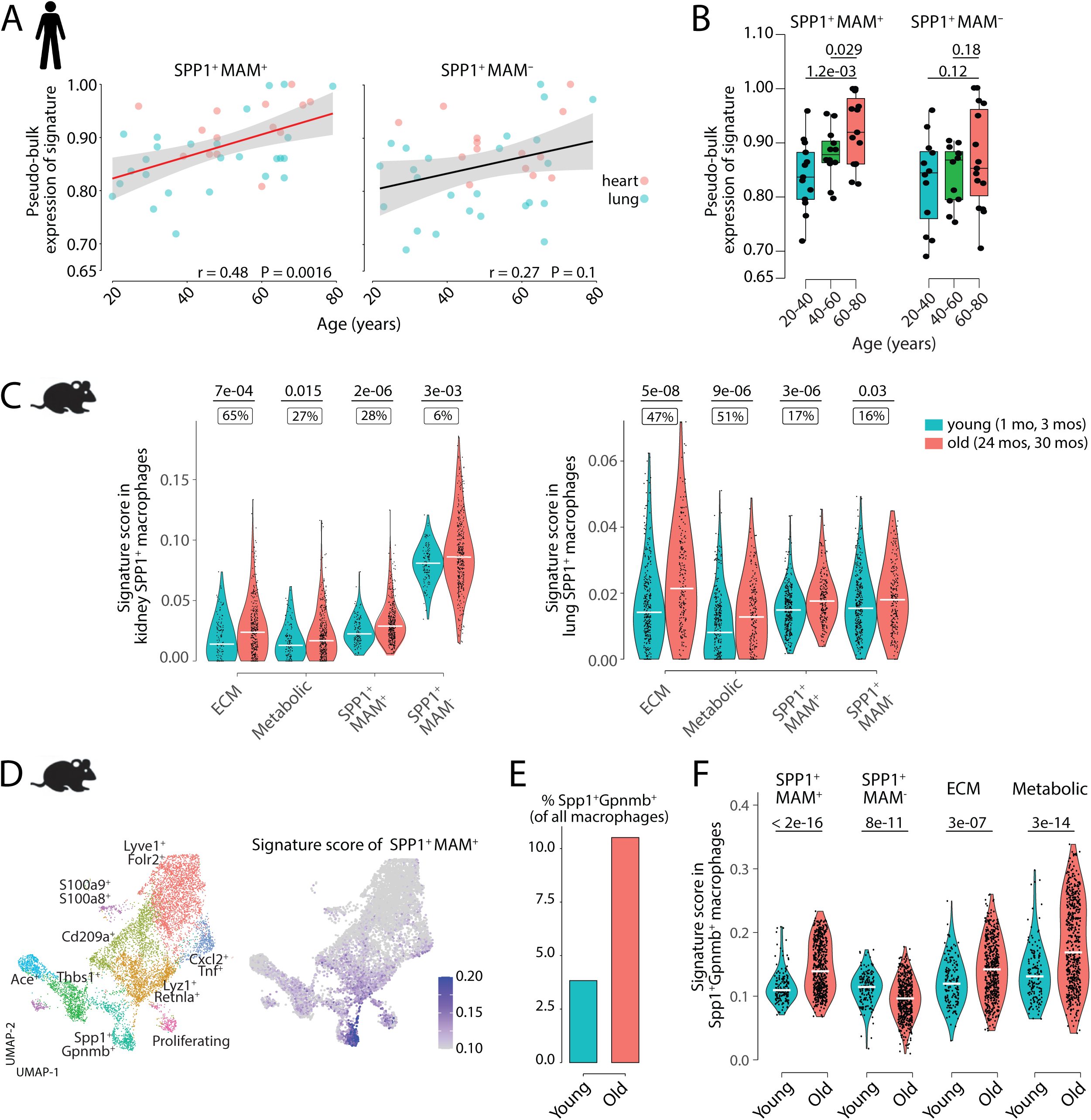
SPP1+MAM+ are associated with aging. (**A)** Scatterplot evaluating the pseudo-bulk expression of signature of SPP1+MAM+ (*left*) or SPP1+MAM- macrophages (*right*) (*y-axis*) from the lung and heart of healthy individuals against age (*x-axis*). Here, for each sample the pseudo-bulk expression of the signature was normalized by the highest expression in each tissue. For each regression, the grey band around the line represents the 95% confidence interval of the regression line. Number of individuals: heart, n = 14; lung, n = 26. (**B**) Boxplot summarizes the pseudo-bulk expression of the signature of SPP1+MAM+ (left) or SPP1+MAM- macrophages (*right*), stratified by age groups (20-40, 40-60 and 60-80 years old). (**C**) Violin plots summarizing the transcriptomic signature of SPP1+MAM+ ECM, SPP1+MAM+ metabolic, SPP1+MAM+, and SPP1+MAM- macrophages in SPP1+ macrophages from the kidney (*left*) and lungs of aged (24-30 months) or young mice (1-3 months). The % in boxes indicate the relative increase in the cell proportion from young to old mice. Kidney: (aged) n = 385 cells, (young) n = 131 cells; lung: (aged) n = 209 cells, (young) n = 339 cells. (**D**) UMAP dimensionality reduction plot of skeletal muscle macrophages from aged (23 months) and young mice (3 months) (n = 3 in each group), where each macrophage is labelled by the cell-type annotation as in the original publication [48] (*left*). Number of cells in each cluster: *Lyve1+Folr2+*, n = 2,962; *Thbs1+*, n = 981; *Lyz1+Retnla+*, n = 1,680; *Cxcl2+Tnf+*, n = 494; *Ace+*, n = 523; *S100a9+S100a8+*, n = 271; *Cd209+*, n = 1,570; *SPP1+Gpnmb+*, n = 710. UMAP dimensionality reduction plot of skeletal muscle macrophages each cell is coloured by the signature score of SPP1+MAM+ (*right*). (**E**) Bar plot displaying the percentage of Gpnmb+SPP1+ macrophages out of total number macrophages (*y-axis*) in young or old mice (*x-axis*). (**F**) Violin plots showing the signature score of SPP1+MAM+, SPP1+MAM-, SPP1+MAM+ ECM or SPP1+MAM+ metabolic process (*y-axis*) in Gpnmb+SPP1+ skeletal muscle macrophages from young or old mice (*x-axis*). Unless otherwise indicated, two-tailed Wilcoxon rank- sum test was used to evaluate the statistical significance (P-value) of differences between two groups. Number of cells: old mice, n = 435; young mice, n = 65.

To provide additional replication of these findings, we looked at a separate dataset of macrophages isolated from aging skeletal muscle [48] (Supplementary Figure 14A). Here, SPP1+Gpnmb+ macrophage cluster showed the highest similarity with our SPP1+MAM+ (Figure 5D and Supplementary Figure 14B), and, in keeping with [48], this cluster is ∼3-fold more represented in old than in young mice (Figure 5E). The expression level of the SPP1+MAM+, SPP1+MAM+ ECM and SPP1+MAM+ metabolic signatures are increased in old Gpnmb+SPP1+ compared with young mouse macrophages (Figure 5F). Opposite to the SPP1+MAM+ signatures, the SPP1+MAM- signature score was significantly decreased in old compared with young mice, suggesting the association with increased aging is specific to the activated SPP1+MAM+ polarization state.

## DISCUSSION

A plethora of studies suggested a specific transcriptional profile that is present in tissue resident macrophages - not only during health and disease in general, but also during pathologies characterized by organ fibrosis. Remarkably, this macrophage state is broadly characterized by the expression of lipid metabolism-associated genes. Depending on the tissue or disease-context, they have been described as LAMs (obesity, [36]) or LAM-like (homeostasis, [15]) or TAMs (cancer, [35]) or DAM (Alzheimer’s disease [37, 38]) or lipid-droplet-accumulating microglia (LDAM, aging brain [49]). Inarguably many genes that characterize these macrophage activation states show context-specificity; nonetheless, certain genes have been repeatedly found throughout different studies and they include *TREM2*, *FABP5*, genes belonging to cholesterol metabolism (*APOE*, *APOC1*, *LPL*) and cell adhesion (*SPP1*, *CD9*). This ubiquitous macrophage transcriptional state seems to be conserved across tissues and is seen during homeostasis and disease [14-16, 20, 22, 24, 25, 27, 34-38, 50]. Scar associated macrophages (SAMs) also share lipid-associated transcriptional markers and although they associate with hepatic and pulmonary fibrotic disease [24, 25, 50], their phenotypic difference with regard to the ubiquitous lipid-associated macrophage state remain incompletely understood. In this study, we focus on SPP1+ macrophages and extend their known association with lung and liver fibrosis to other tissues such as the endometrium. Interestingly, fibrosis is an increasingly recognized feature of endometriosis [12, 51]. Given the recent cell atlas of endometrium revealing disease-associated macrophages [52], our results implicate a potential role of SPP1+ macrophages during endometrial fibrosis.

Here we propose osteopontin (*SPP1*) as a suitable marker for pro-fibrotic macrophages in multiple tissues. *TREM2* is another candidate marker gene that has been proposed for such association. In addition to studies performed in humans [25], single cell-based transcriptome analysis in murine models of human fibrotic disease have confirmed the existence of a Trem2 positive macrophage population associated with disease [50, 53–58], even though some studies suggest a pro-resolution role of Trem2 itself [54, 59] or of Trem2+ macrophages [60]. Furthermore, regenerative Trem2+ macrophages mitigate fibrosis after skin transplantation in humans [61]. These studies suggest that Trem2 is not an optimal marker for SAMs, and in line with the proposed overall pro-regenerative role of Trem2+ LAMs [62], Trem2+ macrophages may represent a hybrid state with features of both resident macrophages and infiltrating monocytes during tissue regeneration [25]. This may explain the contradictory results regarding the previously reported pro-fibrotic and pro-resolution properties of Trem2+ macrophages. SPP1, on the other hand, is a biomarker of fibrosis in NASH [63] and interstitial lung disease progression in systemic sclerosis [64]. Unlike Trem2, there is unequivocal evidence for the pro-fibrotic role of SPP1 during multi-organ fibrosis. Importantly, Spp1 deletion or neutralization in mice attenuates fibrosis in different models of kidney, heart, lung, liver, skin, prostate and muscle injuries [65–76]. Although SPP1 has also been described to be localized to non-myeloid cells [77], macrophage-derived SPP1 can induce migration and proliferation of fibroblasts [64].

Despite being a marker of SAMs, *SPP1* has also been described as a marker of the generalized lipid-associated macrophage state. Lipid-related pathways, and in particular, fatty acid oxidation (FAO) in non-myeloid cells has been previously linked to renal fibrosis [78] and targeting macrophage FAO holds therapeutic potential in pulmonary fibrosis [79]. To gain insights in the specificity of these tissue macrophage functions beyond lipid-related pathways, here we provide a detailed single cell map of the transcriptional heterogeneity of SPP1+ macrophages, including the identification of a matrisome-associated macrophage polarization state. This activation state is characterized by the up-regulation of genes belonging to ECM remodeling such as MMPs, TIMPs and cathepsin family genes. We propose that this metabolically-demanding degradative macrophage state is the result of failure of the ongoing resolution of inflammatory responses and is conserved across tissues. Fibrosis is a non-resolving wound healing process and macrophage infiltration coincides with expression of matrisome-related genes within the skin wound bed [80]. While the general view is that collagen deposition is solely attributed to myofibroblasts, recent reports implicate macrophages as direct contributors to fibrosis during heart repair [81]. Hence the contribution of macrophages toward multi-organ fibrosis could be two-fold; through cell-cell interaction with fibroblasts (as shown in different tissues [6, 82–88]) and/or via cell autonomous production of ECM proteins such as collagens [81]. Therefore, it is tempting to speculate that the transition from SPP1+ to the MAM polarization state corresponds to a ‘shifting gear’ in macrophage whereby the fine balance between resolution and fibrosis is in favour of the latter. Interestingly, SPP1+MAM- macrophages are enriched for secretory (chemokine-chemokine receptor pathways) suggestive of cell-cell interaction, while SPP1+MAM+ are enriched for expression in matrix metalloproteinases (MMP7 and MMP9) previously described to be targets in pulmonary fibrosis [89]. Future work is required to optimize the tissue sorting of MAMs for precise metabolism and proteome profiling, as well as their *in situ* characterization based on markers specifically induced in this cell state (e.g., *MMP7*). The transcriptomic signature of SPP1+MAM+ macrophages can also be used to study the effect of naturally occurring genetic variation on fibrotic disease – similar approaches were proposed for macrophage-driven inflammatory disease [90]. As per the MAM nomenclature, we refer to this as a state of polarization rather than a macrophage subpopulation and make systematic distinction between SPP1+MAM- and SPP1+MAM+. As pertinently argued recently [62], and in accordance with findings presented here, SPP1 expressing LAMs are found in steady state, thus nomenclature based on activation trajectories of these cells (rather than pathology-related states) could be more appropriate. Exploring common and tissue-specific signals that regulate LAMs, including their enhancer landscapes [91] will define these cells with better accuracy.

Mechanistically, our analyses suggest NFATC1 and HIVEP3 as transcription factors specifically associated with the SPP1+MAM+ state, while other TFs (e.g., THRA, NR1H3, ETV5 and MAF) had continuous regulon activity throughout the differentiation of macrophages. This suggests that NFATC1 and HIVEP3 are ‘MAM polarization regulons’ while the other regulons control a wider differentiation trajectory, spanning from infiltrating monocytes to the MAM state. NFATC1 is a well-known master regulatory transcription factor in osteoclast differentiation [92]. Interestingly, HIVEP3 (also known as KRC or ZAS3) is a regulator of osteoclast/osteoblast differentiation and multinucleated giant cell formation [93–95], and an activator of NFATC1 in osteoclasts [95]. These results suggest a concerted action of HIVEP3 and NFATC1 in regulating MAM genes among which some are highly active in osteoclasts (e.g., *MMP9* [96]). The osteoclast resemblance of MAMs may also explain their ECM remodelling concomitant with increased oxidative phosphorylation. Osteoclasts depend on mitochondrial oxidative phosphorylation for their resorptive activity that remodels the bone matrix [97, 98]. As the name suggests, osteopontin was first cloned in bone [99, 100] and has been proposed to mediate the adhesion of osteoclasts to resorbing bone [101, 102]. SPP1 is a constituent of ECM but can also be found as a secreted soluble factor. In ECM-rich fibrotic tissues, SPP1 can mediate the kinetics of the attachment of the macrophages to the evolving extracellular microenvironment. Across tissues and different fibrotic conditions, we found SPP1 as the most significantly and consistently up-regulated transcript associated with the SPP1+MAM+ state, which corroborates this hypothesis. Interestingly, Trem2+ LAMs expressing Spp1 showed a transcriptional profile similar to osteoclasts during advanced stages of murine atherosclerosis [103], and multinucleated adipose tissue macrophages, probably derived from SPP1-expressing LAMs as a result of prolonged obesity, also show functional and morphological resemblance with osteoclasts [104].

A converging body of evidence now supports the infiltrating monocyte origin of pro-fibrotic tissue macrophages [7, 50, 55, 85, 105]. Our trajectory analysis extends this observation to six different human tissues, adding further resolution to the SPP1+ macrophages. According to the continuum model of monocyte-to-macrophage differentiation (or activation paths [106]), a healthy tissue environment is maintained by environmental cues and the ability of the monocyte-derived macrophages to undergo homeostatic differentiation into tissue resident macrophages [107]. The disease state is accompanied by monocyte-derived macrophages infiltrating tissues, and progressively losing the ability to support tissue resident macrophages because of the pathologically evolving environmental cues [107]. For instance, SPP1+ macrophage recruitment to the fatty liver from circulating monocytes coincides with the lack of Kupffer cells [50]. Hence the fine balance between tissue repair and fibrosis, could be explained by these spatiotemporally regulated infiltrating monocytes and their differentiation state (homeostatic/repair *vs* inflammatory/pro-fibrotic) within the tissue microenvironment [107]. Our results fit with this model. We propose that the MAM+ state is an advanced polarization state of SPP1+ macrophages that have been exposed to prolonged inflammatory and fibrotic environmental cues in different tissues. Failing to undergo homeostatic repair function, monocyte-derived SPP1+ macrophages could further polarize towards the SPP1+MAM+ state.

Aging is characterized by low grade inflammation, and here we report an association of MAMs in aged heart and lung in humans, which we replicated using separate murine models of aging. Although low in frequency, we report the SPP1+MAM+ state of activation also in homeostatic tissues. This observation is in line with SPP1+ LAM-like cells identified in healthy human organs [15] including healthy human liver, in proximity to the bile ducts (bile duct LAMs or BD-LAMs [62, 108]). In healthy mice kidney, lung and skeletal muscle tissues, we report SPP1+MAM+ macrophages, and to a lower extent SPP1+MAM-, as significantly increased in old compared with young animals. This may suggest that some markers of SPP1+MAM+ are biomarkers of aging. One of the hallmarks of aging is cellular senescence and SPP1+ DAM have been reported to undergo senescence [109]. Furthermore, senescent Cd4+ T cells robustly upregulate SPP1 at the expense of typical T cell lymphokines [110] and there is augmented macrophage-derived SPP1 in aging skeletal muscle [111]. Since SPP1 is further up-regulated in (and a gene marker of) the MAM state, and given the causal link between senescence and fibrosis [112], our results suggest SPP1+MAM+ cells can hold a senescence-related signature. We found that this is the case in old skeletal muscle macrophages whereby the SPP1+Gpnmb+ senescent macrophages are significantly enriched for the SPP1+MAM+ transcriptional signature. Notably, this enrichment was specific to SPP1+MAM+ state and not found in the general SPP1+ macrophages. In view of the progressive differentiation of SPP1+ macrophages, the MAM state may represent a late stage of differentiation, with the frequency of MAM cells increasing during physiological aging of the tissues that show features of cellular senescence. The differentiation from SPP1+MAM- toward SPP1+MAM+ is accelerated during fibrotic disease, and we speculate the existence of a ‘tipping point’ that can break the balance between tissue regeneration and fibrosis.

Overall, the findings presented in this study indicate an advanced polarization state of monocyte-derived macrophages that show common activation pathways related to multi-organ fibrosis. The age-dependent MAM state may be a result of prolonged inflammatory cues within each tissue microenvironment. A deeper understanding of MAM state across different tissues can refine anti-fibrotic treatment initiatives.

## MATERIALS AND METHODS

### Derivation of monocytes and macrophages clusters from each dataset

Raw counts or already pre-processed expression matrices for the datasets used in this study (Table 1) were downloaded from GEO. R package Seurat (Seurat 4.2.0) was used to perform subsequent single-cell analysis [113]. Quality control to remove low-quality cells and lowly-expressed genes was first applied to all datasets for integrated analyses. For each dataset, cells expressing between 300 and 5000 genes, less than 15% mitochondrial reads (*MT-*), less than 30% ribosomal reads (*RPS*, *RPL*), and less than 0.1% haemoglobin reads (*HBA*, *HBB*) were retained and the rest were filtered out. Only genes expressed in more than 10 cells were retained for the downstream analysis. Raw counts for each cell were log-normalized based on library size, and Seurat’s default integration pipeline (https://github.com/satijalab/seurat/blob/master/vignettes/integration_introduction.Rmd) was applied to integrate each patient sample. Integrated data was scaled, with the number of expressed genes and % of mitochondrial reads in each cell regressed out. Scaled matrix was then subjected to principal component analysis (PCA) on top 2,000 highly variable genes, and the optimal number of PCs for further dimensionality reduction and construction of the nearest neighbor graph was determined by the Elbow method. Briefly, the percentage variance explained by each PC was plotted, and the number of PCs was chosen at the point where a substantial drop in percentage variance occurred. Subsequently, Louvain cluster analysis was performed with Seurat’s FindCluster function with cluster resolution set at 0.5. Analysis of differentially expressed genes (DEG) was performed for each cell cluster using Seurat’s FindAllMarkers function with default parameters. *PTPRC+ CD68+* clusters (putative myeloid cells) expressing monocyte/macrophage related DEGs (see Supplementary Table 1) were extracted for further analysis. Putative myeloid cells were re-normalized, integrated, and clustered as described above. Cell clusters with DEGs containing markers of NK cells (*GZMB, GNLY, CCR7*) [35], dendritic cells (*CD1C, FCER1A, CLEC10A*) [35], non-classical monocytes (*LST1, CDKN1C, FCGR3A*) [35] stressed cells (*HSPA1/A1B, DNJAB*) [114] proliferating cells (*STMN1, TUBB, TOP2A*) [35], and mesenchymal contaminants (*DCN, LUM, COL1A1*) [115] were filtered out to derive the final set of macrophage/monocytes in the dataset, comprising 22,459, 120,273, 8,119, 2,462, 6,739, and 6,839 macrophages from the heart, lung, kidney, skin, liver, and uterus endometrium, respectively.

### Integration and clustering of monocytes/macrophages at the tissue level

The expression matrices retained from the previous step were further subset for genes expressed in all tissues (11,117 genes). Gene expression matrices from cells in the same tissue are aggregated, normalized, and integrated. Integrated count matrix was scaled and subjected to PCA. To derive sub- clusters within the macrophage population in an unbiased manner, Louvain cluster analysis was performed using Seurat’s FindCluster function. Specifically, the analysis was performed multiple times with different cluster resolutions (from .2 to 1 in increments of .05), and the cluster result at each resolution was assessed by the Silhouette score [116]. Briefly, Silhouette score evaluates the goodness of clustering by measuring for every cell the similarity with cells in the same cluster as compared to the similarity to cells in other clusters. We considered a cluster resolution of 0.2 (or lower) as “under-clustering”, and in each case, we selected a resolution that optimizes Silhouette score (Supplementary Figure 1–6A). We note that this approach, though completely data-driven, may produce numerous clusters that share little transcriptomic differences. To merge clusters with small transcriptional differences, we constructed a pseudo-bulk expression profile of each cluster and assessed transcriptional similarity between every cluster-pair using the Spearman correlation of the pseudo-bulk expression profiles of the first 2,000 highly variable genes in the dataset (Supplementary Figure 1–6B). To that aim, we employed the method described by Shafer et al. [117]. For every expressed gene in a dataset, the pseudo-bulk expression of the gene in a cluster is normalized by the sum of its pseudo- bulk expression of all the clusters in the dataset. The normalized expressions of the top highly variable genes are then used for calculation of Spearman correlation. Cluster pairs with Spearman correlation greater than 0.6 were merged (Supplementary Figure 1–6C). DEGs for the merged clusters were calculated by Seurat’s FindAllMarkers function. Each cluster was annotated by the top DEGs in reference to known marker of homeostatic macrophages and monocytes, as well as *TREM2+* macrophages (which have been implicated in multiple human diseases such as cancer [35], obesity [36], atherosclerosis [118], and neurodegenerative diseases [119] (see Supplementary Table 1, Supplementary Table2) (Supplementary Figure 1–6D). We plotted the top 15 DEG with the highest fold change for each cluster in a heatmap in Supplementary Figure 1–6E, and highlighted the above-mentioned known markers.

### Identification of a matrisome-associated macrophage (MAM) polarization state

The standard Seurat integration pipeline (as mentioned above) was used to integrate SPP1+ macrophages from all organs (lung, liver, heart, skin, endometrium, and kidney), using the top 3,000 highly variable genes as feature genes in the integration. This resulted in pooling 2,454, 8,169, 1,113, 153, 792, and 1,438 SPP1+ macrophages from the heart, lung, kidney, skin, liver, and uterus endometrium, respectively. To reduce the representation of SPP1+ lung macrophages in the combined analysis, SPP1+ macrophages from the Adams et al IPF dataset (n = 13,362) were excluded from this analysis. Integrated matrix was scaled, subjected to PCA, and clustered as described above (see Supplementary Figure 7A-C). Three clusters were identified at resolution 0.25 (Supplementary Figure 7), and GSEA analysis was further performed on the DEGs in each cluster, querying the MSigDB Human C2 gene-set database [120]. Cluster 2 was enriched for inflammatory process (e.g., “TNFA Signaling Via NF-KB”, “Inflammatory response”) (Supplementary Figure 7D), and cluster 0 homeostatic process (e.g., “Ribosome”, “Eukaryotic Translation and Elongation”) (Supplementary Figure 7E). Cluster 1 was enriched for ECM and metabolic processes, and denoted as “matrisome-associated macrophage” and referred to as SPP1+MAM+ (see SPP1+MAM+ transcriptomic signature for details).

### SPP1+MAM+ transcriptomic signature

We derived three transcriptomic signatures of SPP1+MAM+, a core SPP1+MAM+ signature (284 genes, Supplementary Table 4), an SPP1+MAM+ ECM-related signature, (42 genes, Supplementary Table 6) and a SPP1+MAM+ metabolism-related signature (53 genes, Supplementary Table 7). To construct the core signature of SPP1+MAM+, we combined two criteria: (1) change in expression (across tissues) and (2) number of SPP1+MAM+ cells expressing a given marker gene. First, fold changes in expression between SPP1+MAM+ and SPP1+MAM- macrophages were retrieved using Seurat’s FindMarkers function (logfc.threshold = -Inf, min.pct = -Inf, min.diff.pct = -Inf), which was run separately for each tissue. Then, the fold changes (FC) of each gene (the higher the log2FC the more differentially expressed in SPP1+MAM+) were then averaged across tissues. Second, we calculated for each gene the specificity of expression for the gene to SPP1+MAM+ using the difference in proportions of cells expressing a gene between SPP1+MAM+ and SPP1+MAM- macrophages. A one-tailed z-test was then performed to determine the statistically significant difference between cell proportions (i.e., proportion SPP1+MAM+ - proportion SPP1+MAM- macrophages). Genes were selected according to both criteria: (1) averaged cross-tissue log2FC >0.25 and (2) alpha ≤ 0.05 (z-test), corresponding to a difference in cell proportion between SPP1+MAM+ and SPP1+MAM- ranging from 0.091 to 0.4.

To derive the ECM or metabolism-related signature, GSEA analysis was performed on the averaged log2FC, using the R package ClusterProfiler (clusterProfiler 3.18.1) [121]. We queried the MSigDB Human C2 gene-set database [120] and collected the ECM and metabolism-related pathways in the top 100 annotation terms that were significantly enriched in SPP1+MAM+ (FDR < 0.05, NES > 1.6). We then pooled together the core enrichment genes (leading edges) contributing to the enrichment of the extracted processes. The ECM-related signature was finally derived by intersecting the core enrichment of ECM-related pathways and the core SPP1+MAM+ signature, and the metabolism-related signature was constructed in the same manner.

### SPP1+ MAM+ gene markers

COMET (COMETSC 0.1.13) was used to derive gene markers of SPP1+MAM+ macrophages [40]. Briefly, COMET is a marker-panel selection tool for single-cell data, which employs XL-minimal Hypergeometric test to binarize the expression of a gene and assess the extent the gene could be a good marker for a cell cluster. The whole expression matrix of the SPP1+ macrophages (SPP1+MAM+ and SPP1+MAM-) and the cluster assignment were used as input to COMET, which was run using default settings for 4- marker panels. We noted that the same genes occurred multiple times in 1,366 unique combinations of 4 distinct gene markers (marker panels) predicted by COMET with P=9x10-136. We therefore considered all 4-markers panel predictions, ranked each gene based on the number of occurrences in the prediction and selected the top 6 genes as marker genes that distinguish SPP1+MAM+ from SPP1+ macrophages using the Elbow method (Supplementary Figure 8A). Briefly, on a scatterplot we ranked each marker gene from COMET based on the number of occurrences, with the x-axis the rank and the y-axis the number of occurrences. We drew a line between the dots representing the top ranked gene (i.e., SPP1) to the least ranked gene, and calculated the shortest distance of each dot (which represents a marker gene) to the line. We derived the marker gene that maximizes this distance, and retained this gene and other genes ranked higher as markers of SPP1+MAM+ macrophages. UMAP of the 6 SPP1+MAM+ marker genes predicted by COMET are reported in Figure 2B.

### Comparison of SPP1+MAM+ transcriptomic signatures with LAMs, TAMs and DAM

Expression level of ECM and metabolic signatures in SPP1+MAM+ was compared to tumor-associated macrophages (TAMs), lipid-associated macrophage (LAMs), and disease-associated microglia (DAM). Expression data of TAMs from Mulder et al [35] was obtained from https://gustaveroussy.github.io/FG-Lab/; LAMs from Jaintin et al [36] and Wirka et al [118] were obtained from GEO GSE128518 and GSE131780 [36, 118]; DAM from Kumar et al obtained from https://epicimmuneatlas.org/NatNeu2022/ [119]. *TREM2+* TAM (“cluster 3 macrophages” in Mulder et al. [35]) from lungs, liver, skin, and colon were extracted based on single-cell meta information provided. The adipose tissue LAMs from Jaintin et al [36] were extracted also based on the provided cell-type annotation. To extract LAMs (from coronary atherosclerotic plaque), raw expression data from Wirka et al. was processed following the authors’ methods as closely as possible, using the parameters presided by the authors in the original publication [118]. DAM were defined as microglia clusters isolated from temporal lobe epilepsy patients provided by Kumar et al. [119], and only temporal lobe microglia samples processed within the same batch were kept (‘P6.A’, ‘P4’, ‘P5.A’, P3.A’). The microglia were then scored for transcriptomic signature of DAM using Seurat’s AddModuleScore function using default parameters (see Supplementary Table 2 for marker genes of this signature), and a z-score of DAM signature score was calculated. *TREM2+APOE+SPP1+* microglia clusters with the highest median signature score of DAM, i.e., in the top 25th percentile of z-score (corresponding to DAM score = 0.349) were retained as DAM for further analysis (Supplementary Figure 9). In order to compare expression levels in tissue macrophages derived from different tissues, SPP1+MAM+, DAM, LAMs, and TAMs were integrated using Seurat’s integration pipeline. First, SPP1+MAM+ and TAMs were split into smaller datasets by tissue, and the expression data of each tissue-specific SPP1+MAM+ and TAM, as well as LAM and DAM, were normalized by SCTransform with default parameters. To integrate expression data of different macrophages, we used Seurat’s reference-based integration pipeline for data normalized with SCTransform and selected the top 3,000 highly variable genes in the datasets as feature genes. SPP1+MAM+ (separated by tissue and normalized separately) were used as the reference for other macrophages to be projected onto. The R package UCell (UCell 1.3.1) was used to evaluate the transcriptomic signature of SPP1+MAM+ ECM and SPP1+MAM+ metabolism in the pooled macrophage dataset using integrated count matrix [122]. Briefly, UCell calculates gene signature score based on Mann-Whitney U statistics; the algorithm relies on relative gene expression in each individual cell, instead of on dataset composition, which further mitigates batch effect and allows unbiased cross-tissue/dataset comparison of transcriptomic signatures [122].

### Trajectory and cell differentiation propensity analyses

#### Differentiation trajectory analysis

The R package Slingshot (Slingshot 1.4.0) was used to perform differentiation trajectory analysis on macrophages, separately in each tissue [123]. Briefly, Slingshot derives differentiation paths from a specified origin and calculates for each cell a pseudotime, which approximates differentiation progression of a cell toward the destination of the trajectory. The UMAP dimensionality reduction and cell-type labels (e.g., *FCN1^+^* monocyte, *MARCO+* macrophages, *etc*.) were used as input, and *FCN1^+^* monocyte was specified as the origin of differentiation. One trajectory leading to SPP1+MAM+ and the others to homeostatic macrophage (*RNASE1^+^*, *FABP4+*, and *MARCO+* macrophages) were consistently recapitulated across the different tissues. We further applied Monocle (Monocle 3.16) [124] to validate the differentiation trajectories predicted by Slingshot. Monocle was run with default setting using UMAP dimensionality reduction and cell-type labels as input.

#### Differentiation propensity analysis

We further extracted SPP1+MAM+, transitional macrophages, and SPP1+MAM- macrophages to examine the propensity of a SPP1+MAM- macrophage to differentiate into SPP1+MAM+ macrophage in disease or control. Diffusion map was employed to derive the probability of a given cell to differentiate into SPP1+MAM+. The diffusion map is a dimensionality-reduction algorithm that recovers distance measure between a pair of cells with respect to the transitional probability from one cell to another based on random walk. We approximated the differentiation potential of a cell using transitional probability. The R package Destiny (destiny 3.10.0) was used to build a diffusion map on transitional macrophages, SPP1+MAM- macrophages and SPP1+MAM+ separately for each tissue [125]. Expression data of the top 2,000 highly variable genes was used as input. The propensity of SPP1+MAM- macrophages to differentiate into SPP1+MAM+ was approximated as the quotient of the probability of a cell to transition into SPP1+MAM+ over the sum of the probability of SPP1+MAM- to transition into SPP1+MAM+ or transitional macrophage. Since we focus on assessing probability of cell transitions, we are not considering the probability of cells retaining the same state.

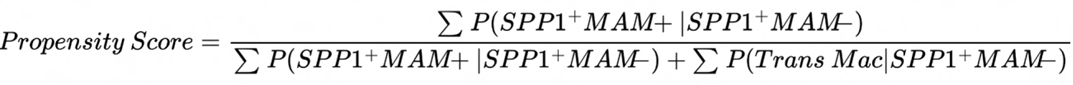

where P(MAM|SPP1+) denotes the probability of a SPP1+MAM- macrophage to transition into SPP1+MAM+, and P(Trans Mac|SPP1+) denotes the probability of a SPP1+MAM- macrophage to transition back to transitional macrophage.

### Regulon analysis

Regulon (gene-regulatory network) analysis was performed using pySCENIC (version 0.12.0) to derive a set of regulons likely driving the differentiation from SPP1+MAM- to SPP1+MAM+ macrophages [126]. Briefly, pySCENIC (1) derives a set of gene co-expression network defined by a transcription factor (TF) and its target genes, (2) evaluates a network for enrichment of TF-specific cis-regulatory element and removes targets genes lacking an enrichment of these elements, and (3) assesses the activity level of the network in each individual cell by an “Area Under the Curve Score” (AUC). We ran pySCENIC with default parameterization on SPP1+MAM- and SPP1+MAM+ macrophages. The expression data from integrated assay (output of Seurat’s integration pipeline) (13,402 cells by 3,000 genes) were used as input, and a list of human-specific TF was downloaded from <u>github.com/aertslab/pySCENIC/blob/master/resources/hs_hgnc_tfs.txt</u>. For each gene in the transcriptome, a tree-based regression model was built with the TF candidates as predictors using GRNBoost2. In step 2 (network refinement), we used the following database of genome-wide regulatory features (<u>hg19-500bp-upstream-10species.mc9nr.genes_vs_motifs.rankings.feather</u>, <u>hg19-tss-</u> <u>centered-5kb-10species.mc9nr.genes_vs_motifs.rankings.feather</u>) and TF motifs (<u>motifs-v9-nr.hgnc-</u> <u>m0.001-o0.0.tbl</u>) provided by laboratory of Serin Aerts to assess a regulon for the enrichment of regulatory features and prune the member genes. In brief, these database files contain pre-computed rankings of genome-wise regulatory features in the target genes. In step 3 (the evaluation of regulon activity), pySCNEIC ranked each gene in the transcriptome of a cell by its expression, and an AUC score evaluates the enrichment of the members in a regulon based on this ranking. The activation status of a regulon in a cell is finally derived by binarizing the AUC score.

In total, 120 regulons were identified, and we only retained regulons which are activated in at least 10% of the cells in at least 4 tissues. To prioritise relevant regulons, we evaluated the specificity of a regulon to SPP1+MAM+ macrophages using three criteria: (1) the specificity to the activation status of the regulon with respect to SPP1+MAM+ macrophages (relative to SPP1+MAM- macrophages), (2) the specificity of the regulon to the core signature of SPP1+MAM+ (see SPP1+MAM+ transcriptomic signature for details) and (3) the potential of the regulon to promote the polarization of SPP1+MAM- macrophages towards the SPP1+MAM+ state.

#### First criterion

We evaluated the specificity of the activation of a regulon in SPP1+MAM+ using the Diagnostic Odd Ratio (DOR) [127]. Briefly, DOR assesses the odds of a positive test in “cases” relative to the odds of a positive test in “controls”. Here we refer to the activation of the regulon in SPP1+MAM+ as “cases” and the activation of the regulon in SPP1+MAM- macrophage as “control”. The DOR for a regulon is calculated using the following formula:

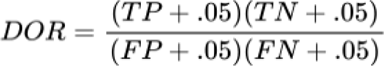

where TP refers to number of SPP1+MAM+ in which the regulon is activated, TN the number of SPP1+MAM- macrophages in which the regulon is not activated, FP the number of SPP1+MAM- macrophages in which the regulon is activated, and FN the number of SPP1+MAM+ macrophages in which the regulon is not activated.

#### Second criterion

to assess the specificity of a regulon to the core signature of SPP1+MAM+ macrophages, we examined (1) the size of overlap between the regulon and the signature and (2) the importance of the genes in the overlap. Specifically, we first performed a hypergeometric test to evaluate the significance of the overlap between the core SPP1+MAM+ signature and member genes of the regulon. We then calculated and summed over the gene expression fold changes between the SPP1+MAM+ to SPP1+MAM- macrophage states of the overlapping genes. We repeated the same calculation (both hypergeometric test and fold change calculation) for genes downregulated in SPP1+MAM+ compared to SPP1+MAM- (log2FC<-0.5) The specificity of a regulon to core SPP1+MAM+ signature was then calculated using the following formula:

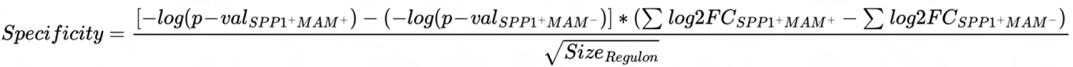

where “p-val” refers to the p-value of the hypergeometric test, and SizeRegulon is the number of genes in the regulon.

#### Third criterion

to delineate which regulon is more specifically required for the polarization of SPP1+MAM- macrophages to the SPP1+MAM+ state (as opposed to the possible differentiation from transitional to SPP1+MAM- macrophages), we pooled together SPP1+MAM+, transitional, and SPP1+MAM- macrophages from all tissue and plotted the expression of a regulon in each macrophage against the expression of SPP1+MAM+ signature (Figure 4C) for selected regulons (see Supplementary Table 9 for member genes of each regulon). Here, we evaluated the potential of SPP1+MAM- macrophages to acquire the SPP1+MAM+ polarization state using the expression of SPP1+MAM+ signature. We performed linear regression analyses of regulon expression against expression of the SPP1+MAM+ signature, using the transitional and SPP1+MAM- macrophages, or using SPP1+MAM- and SPP1+MAM+ macrophages. We classified the regulon with greater increase in regulon expression within the SPP1+MAM- and SPP1+MAM+ macrophages (as compared to within SPP1+MAM- and transitional macrophages) as the regulon most likely to drive the differentiation progression from SPP1+MAM- macrophages to the SPP1+MAM+ polarization state.

### Evaluation of SPP1+MAM+ and SPP1+MAM- macrophage signatures in aging

#### Association of SPP1+MAM+ signature with aging in humans

We extracted human control macrophages of the lungs, heart, and skin from Adams et al. [17], Koenig et al. [20], and Gur et al. [19] dataset respectively (see also Derivation of monocytes and macrophages clusters from each dataset for further details). For the heart control macrophages, we analyzed the single-nuclei RNAseq dataset from Koenig et al. [20] (instead of single-cell RNAseq) as this dataset was larger. Using supplementary data and metadata provided by the authors, we further removed samples labelled as control but with other disease traits, such as non-ischemic cardiomyopathy, heart failure, or abnormal findings on echocardiography at young age. We further removed samples with <10 macrophages from subsequent analysis. UCell was then used to score each healthy tissue macrophage for signature of SPP1+MAM+, SPP1+MAM-, SPP1+MAM+ metabolic process, or SPP1+MAM+ ECM remodeling. Briefly, the “SPP1+MAM- ” signature defined by the upregulated genes in SPP1+MAM- compared to SPP1+MAM+ macrophages, using FDR<0.05, log2(FC) >0.25 (average cross-tissues), and expressed by a significantly higher proportion of cells compared to SPP1+MAM+ by z-test (alpha = 0.05); as described above (see SPP1+MAM+ transcriptomic signature). Pseudo-bulk expressions of these signatures are then constructed per-patient sample by taking the median of the scores. We employed a linear regression model where patient’s SPP1+MAM+ was regressed against the patient’s age, tissue and patient’s gender. To make the pseudo- bulk expression score comparable across different tissues/studies, before combining samples from different tissues/studies in a linear regression modeling, we normalized the per-patient pseudo-bulk expression of the signature by the largest value in each tissue. Age data were available for heart, lungs and skin studies. However, we excluded the skin from this analysis due to the low cell-number per-patient in the skin ([skin]: 5 to 57 cells in each patient sample, [heart]: 311 to 1248 cells, [lung]: 202 to 5928 cells). In addition, skin was removed because in the linear regression analysis of SPP1+MAM+ signature against age, sex, and tissue (combining all samples across skin, heart, and lungs), skin was a significant covariate for the change in SPP1+MAM+ signature (Supplementary Figure 13).

#### Association of SPP1+MAM+ signature with aging in mice

We extracted mouse macrophages from murine single-cell atlas by The Tabula Muris Consortium [47]. Author-provided cell-type annotations were used to extract the macrophages, and we retained the dataset from the young mice (aged 1 ∼ 3 months) or the old (aged 24 ∼ 30 months). Using Seurat standard label transfer pipeline, we unbiasedly projected the mouse macrophages (from tissues present in our dataset, the heart, lungs, kidney, and liver) to our human macrophage atlas, performing this procedure tissue by tissue. We retained for analysis these tissues (the lungs and kidney) where we could detect at least 20 SPP1+ macrophages in both the young and aged group. Each murine SPP1+ macrophage (defined by cross-species projection) was then scored for the signature of SPP1+MAM+, SPP1+MAM-, SPP1+MAM+ ECM and SPP1+MAM+ metabolic process.

We carried out a separate analysis of association between SPP1+MAM+ and aging using sorted macrophages from young and old mice skeletal muscle [48]. For this single cell dataset, we used the R scripts provided by the authors to annotate macrophage clusters [48] (Supplementary Figure 14), and scored macrophage clusters for the signature of SPP1+MAM+, SPP1+MAM-, ECM and metabolic process accordingly. We focused on the SPP1+Gpnmb+ macrophage cluster as this was associated with aging in [48], and was most similar to our SPP1+MAM+ macrophage signature. See Supplementary Figure 14 for details on comparison between SPP1+MAM+ macrophages and the SPP1+Gpnmb+ macrophage cluster [48].

---

**Supplementary Figure 1.**
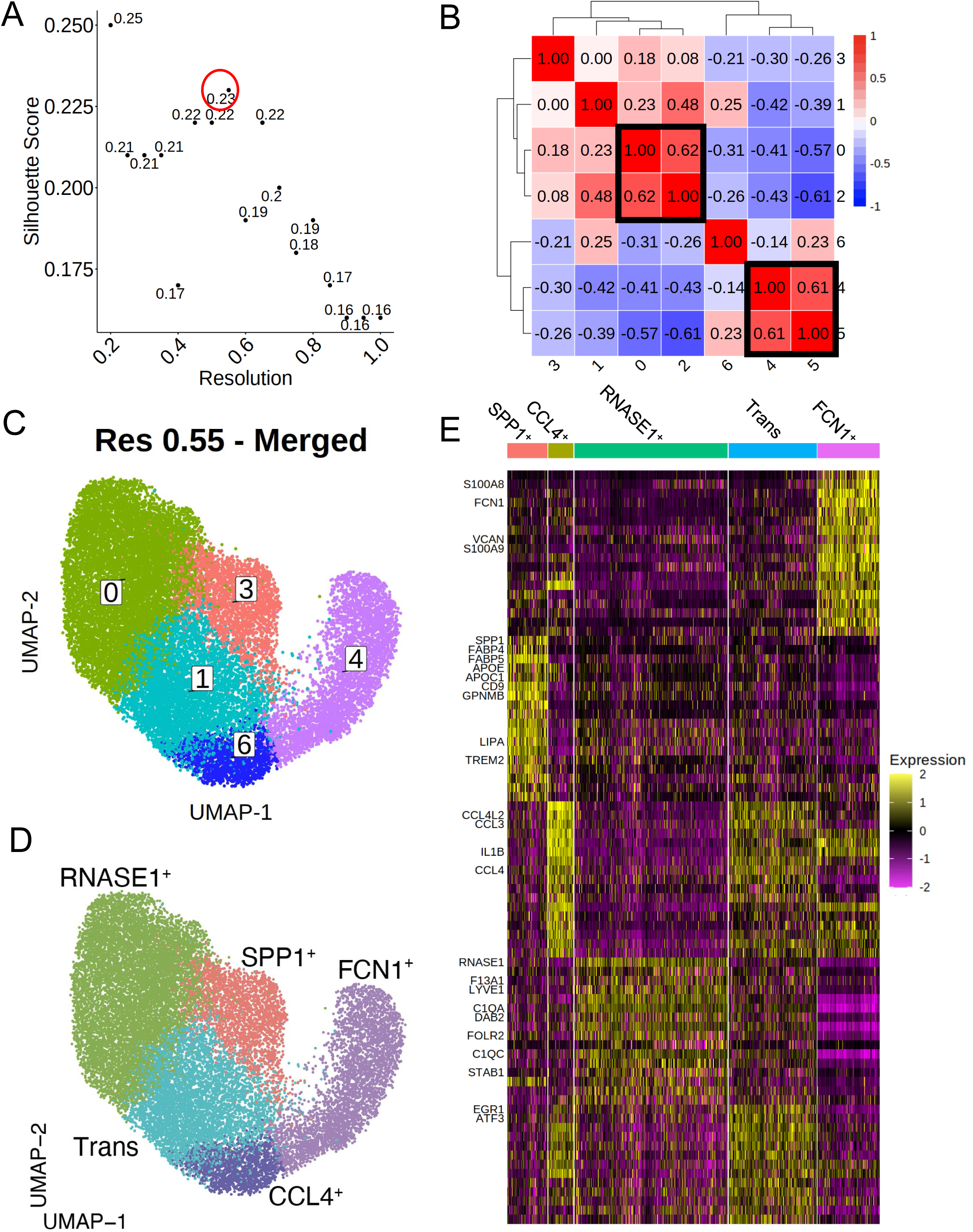
Data-driven clustering of heart macrophages results in five clusters. (**A**) Scatterplot of Silhouette score (see Methods) of single-cell clustering results by Seurat’s Louvain clustering algorithm at different resolutions (“res”). Briefly, res dictates granularity of clustering (i.e., larger res value results in more clusters). A res > 0.2 maximizes the Silhouette score (circled in red); res = 0.55 corresponds to Silhouette score of 0.23. (**B**) Heatmap showing the Spearman correlation between the pseudo-bulk profiles of clusters derived by using the selected res. Each box is labeled by the Spearman correlation between the cluster pair. Clusters with Spearman correlation > 0.6 were merged (cluster pair 4 and 5; cluster pair 0 and 2). (**C**) UMAP projection of heart macrophages, with each cell colored by its cluster assignment. Note that the merged clusters 0 and 2 are labelled as a single cluster 0; the merged clusters 4 and 5 are labelled as a single cluster 4. (**D**) UMAP projection of heart macrophages according to known macrophage markers (see Supplementary Table1) such as FCN1^+^ monocytes, CCL4+ macrophages, transitional (Trans) macrophages, SPP1+ macrophages, and RNASE1^+^ macrophages. (**E**) Heatmap displays the scaled gene expression expressions of top 15 marker genes within each cluster by average log2 fold change. Macrophage markers genes (Supplementary Table 1) and selected marker genes employed for cluster assignment are shown.

**Supplementary Figure 2.**
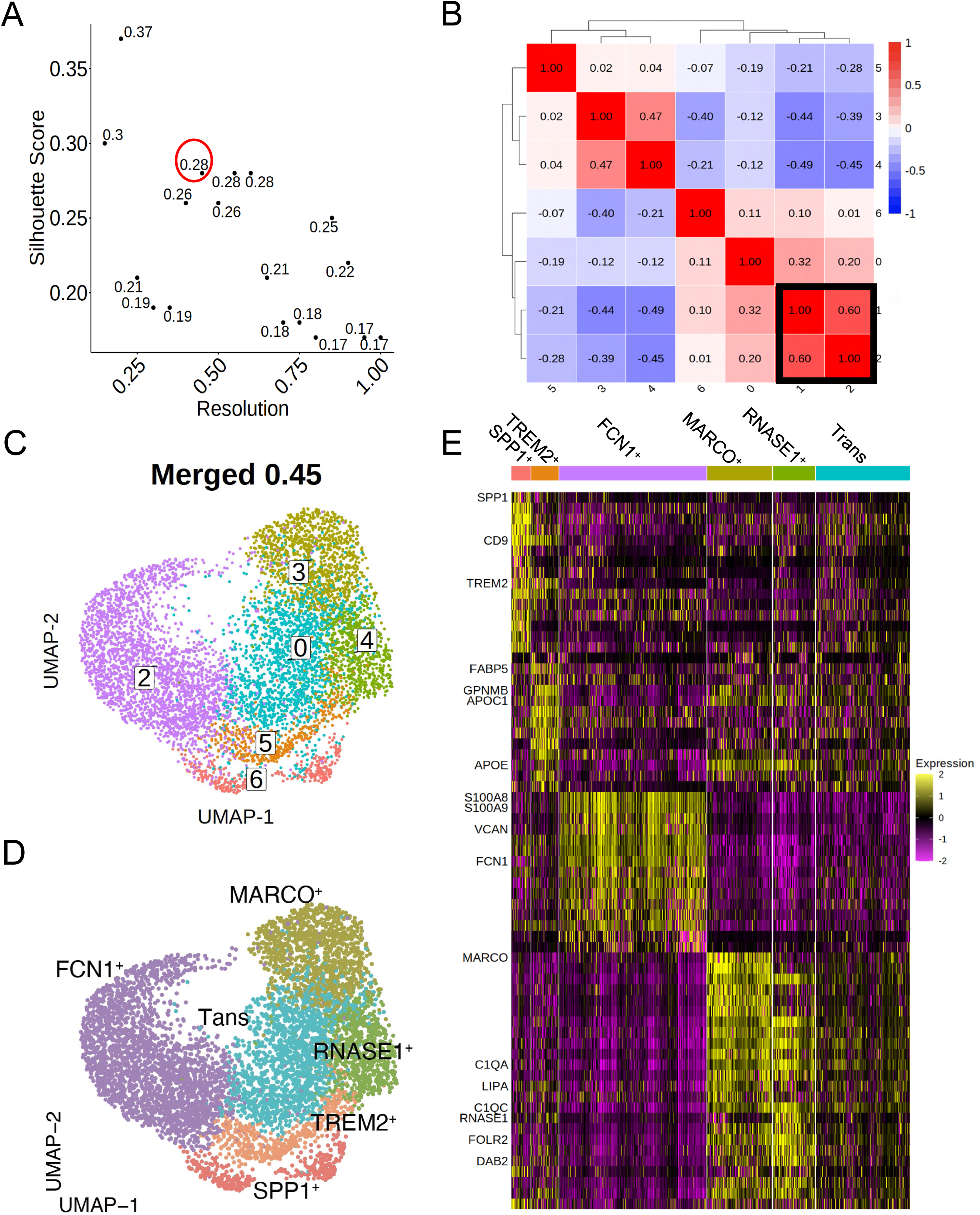
Data-driven clustering of liver macrophages results in six clusters. (**A**) Scatterplot of Silhouette score (see Methods) of single-cell clustering results by Seurat’s Louvain clustering algorithm at different resolutions (“res”). Briefly, res dictates granularity of clustering (i.e., larger res value results in more clusters). A res > 0.2 maximizes the Silhouette score (circled in red); res = 0.45 corresponds to Silhouette score of 0.28. (**B**) Heatmap showing the Spearman correlation between the pseudo-bulk profiles of clusters derived by using the selected res. Each box is labeled by the Spearman correlation between the cluster pair. Clusters with Spearman correlation > 0.6 were merged (cluster pair 1 and 2). (**C**) UMAP projection of liver macrophages, with each cell colored by its cluster assignment. Note that the merged clusters 1 and 2 are labelled as a single cluster 2. (**D**) UMAP projection of liver macrophages according to known macrophage markers (see Supplementary Table 1) such as FCN1^+^ monocytes, MARCO+ macrophages, transitional (Trans) macrophages, SPP1+ macrophages, TREM2+ macrophages, and RNASE1^+^ macrophages. (**E**) Heatmap displays the scaled gene expression expressions of top 15 marker genes within each cluster by average log2 fold change. Macrophage markers genes (Supplementary Table 1) and selected marker genes employed for cluster assignment are shown.

**Supplementary Figure 3.**
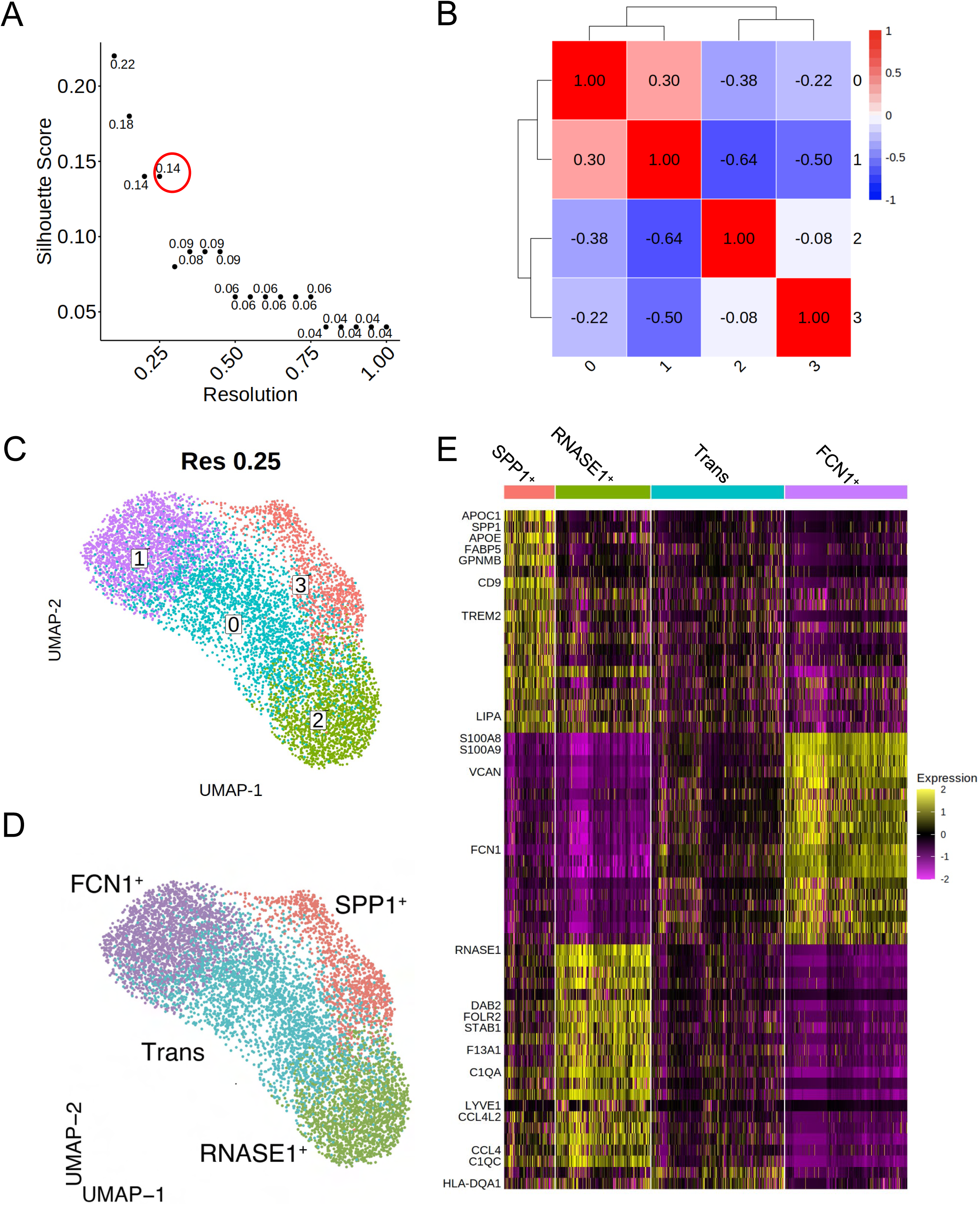
Data-driven clustering of kidney macrophages results in four clusters. (**A**) Scatterplot of Silhouette score (see Methods) of single-cell clustering results by Seurat’s Louvain clustering algorithm at different resolutions (“res”). Briefly, res dictates granularity of clustering (i.e., larger res value results in more clusters). A res > 0.2 maximizes the Silhouette score (circled in red); res = 0.25 corresponds to Silhouette score of 0.14. (**B**) Heatmap showing the Spearman correlation between the pseudo-bulk profiles of clusters derived by using the selected res. Each box is labeled by the Spearman correlation between the cluster pair. (**C**) UMAP projection of kidney macrophages, with each cell colored by its cluster assignment. (**D**) UMAP projection of kidney macrophages according to known macrophage markers (see Supplementary Table 1) such as FCN1^+^ monocytes, transitional (Trans) macrophages, SPP1+ macrophages, and RNASE1^+^ macrophages. (**E**) Heatmap displays the scaled gene expression expressions of top 15 marker genes within each cluster by average log2 fold change. Macrophage markers genes (Supplementary Table 1) and selected marker genes employed for cluster assignment are shown.

**Supplementary Figure 4.**
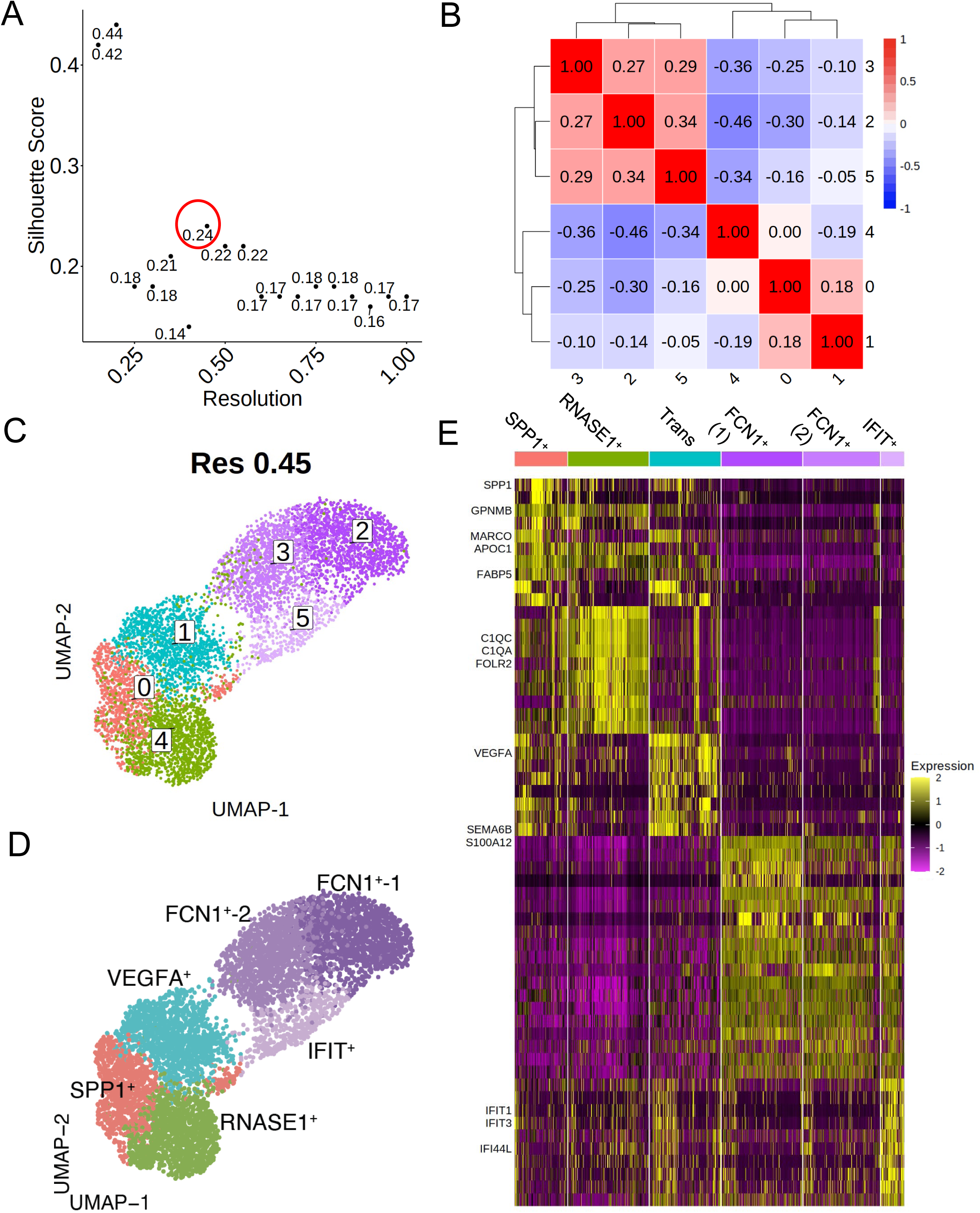
Data-driven clustering of endometrium macrophages results in six macrophage clusters. (**A**) Scatterplot of Silhouette score (see Methods) of single-cell clustering results by Seurat’s Louvain clustering algorithm at different resolutions (“res”). Briefly, res dictates granularity of clustering (i.e., larger res value results in more clusters). A res > 0.2 maximizes the Silhouette score (circled in red); res = 0.45 corresponds to Silhouette score of 0.24. (**B**) Heatmap showing the Spearman correlation between the pseudo-bulk profiles of clusters derived by using the selected res. Each box is labeled by the Spearman correlation between the cluster pair. (**C**) UMAP projection of uterus endometrium macrophages, with each cell colored by its cluster assignment. (**D**) UMAP projection of endometrium macrophages according to known macrophage markers (see Supplementary Table 1) such as FCN1^+^ monocytes, INFIT+ monocytes, VEGFA+ transitional (Trans) macrophages, SPP1+ macrophages, and RNASE1^+^ macrophages. (**E**) Heatmap displays the scaled gene expression expressions of top 15 marker genes within each cluster by average log2 fold change. Macrophage markers genes (Supplementary Table 1) and selected marker genes employed for cluster assignment are shown.

**Supplementary Figure 5.**
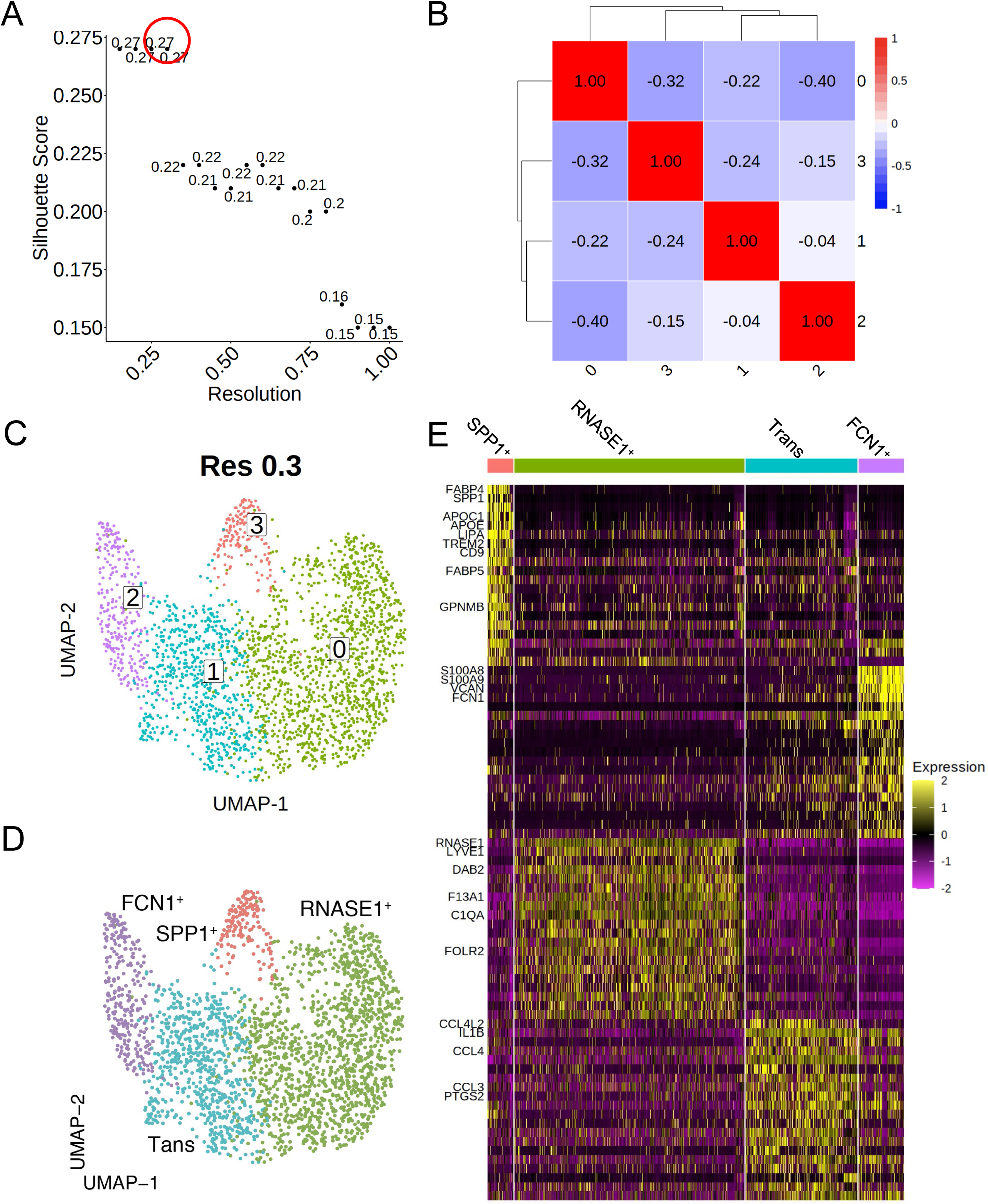
Data-driven clustering of skin macrophages results in four clusters. (**A**) Scatterplot of Silhouette score (see Methods) of single-cell clustering results by Seurat’s Louvain clustering algorithm at different resolutions (“res”). Briefly, res dictates granularity of clustering (i.e., larger res value results in more clusters). A res > 0.2 maximizes the Silhouette score (circled in red); res = 0.3 corresponds to Silhouette score of 0.27. (**B**) Heatmap showing the Spearman correlation between the pseudo-bulk profiles of clusters derived by using the selected res. Each box is labeled by the Spearman correlation between the cluster pair. (**C**) UMAP projection of skin macrophages, with each cell colored by its cluster assignment. (**D**) UMAP projection of skin macrophages according to known macrophage markers (see Supplementary Table 1) such as FCN1^+^ monocytes, transitional (Trans) macrophages, SPP1+ macrophages, and RNASE1^+^ macrophages. (**E**) Heatmap displays the scaled gene expression expressions of top 15 marker genes within each cluster by average log2 fold change. Macrophage markers genes (Supplementary Table 1) and selected marker genes employed for cluster assignment are shown.

**Supplementary Figure 6.**
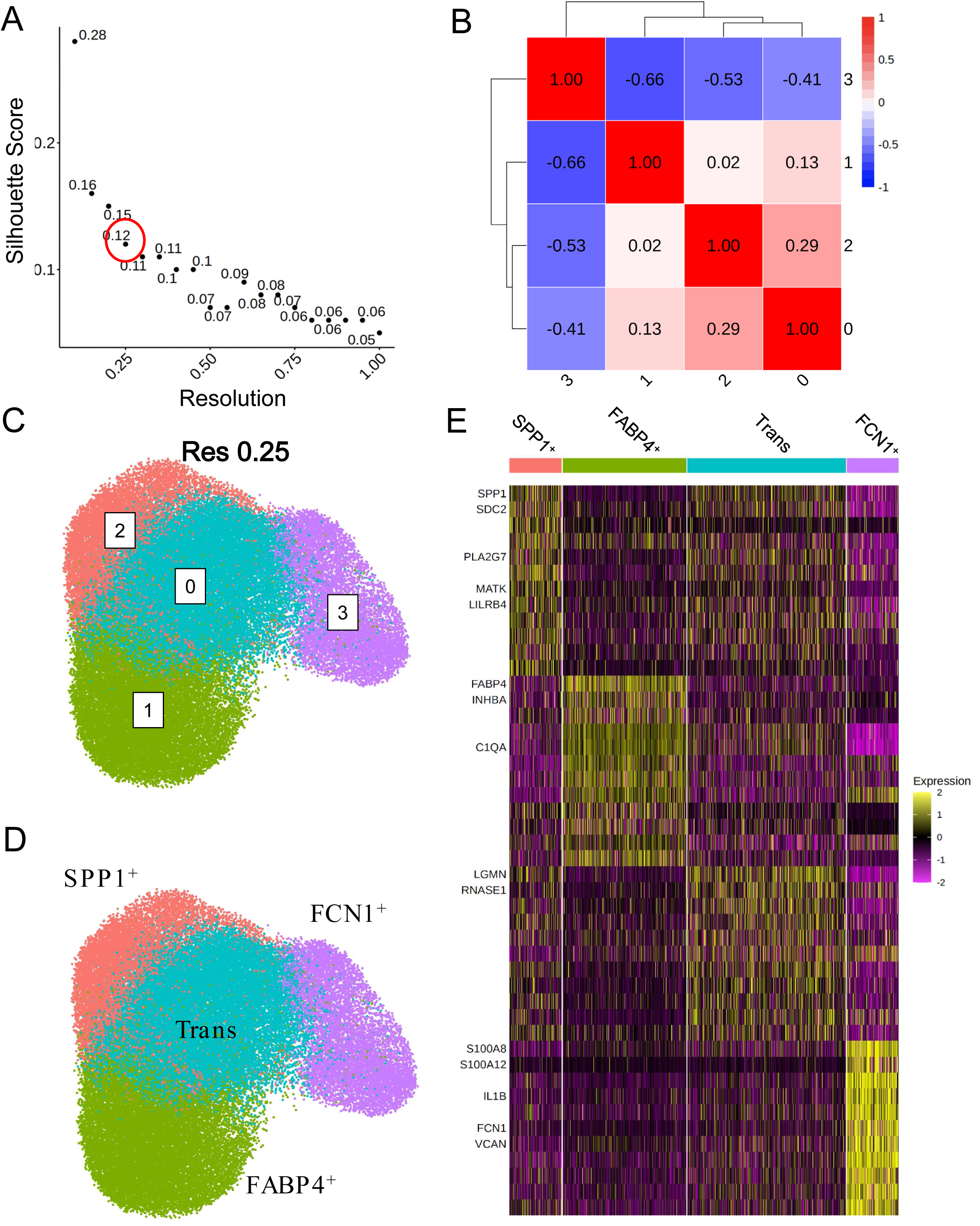
Data-driven clustering of lung macrophages results in four clusters. (**A**) Scatterplot of Silhouette score (see Methods) of single-cell clustering results by Seurat’s Louvain clustering algorithm at different resolutions (“res”). Briefly, res dictates granularity of clustering (i.e., larger res value results in more clusters). A res > 0.2 maximizes the Silhouette score (circled in red); res = 0.25 corresponds to Silhouette score of 0.12. (**B**) Heatmap showing the Spearman correlation between the pseudo-bulk profiles of clusters derived by using the selected res. Each box is labeled by the Spearman correlation between the cluster pair. (**C**) UMAP projection of lung macrophages, with each cell colored by its cluster assignment. (**D**) UMAP projection of lung macrophages according to known macrophage markers (see Supplementary Table 1) such as FCN1^+^ monocytes, transitional (Trans) macrophages, SPP1+ macrophages, and FABP4+ macrophages. (**E**) Heatmap displays the scaled gene expression expressions of top 15 marker genes within each cluster by average log2 fold change. Macrophage markers genes (Supplementary Table 1) and selected marker genes employed for cluster assignment are shown.

**Supplementary Figure 7.**
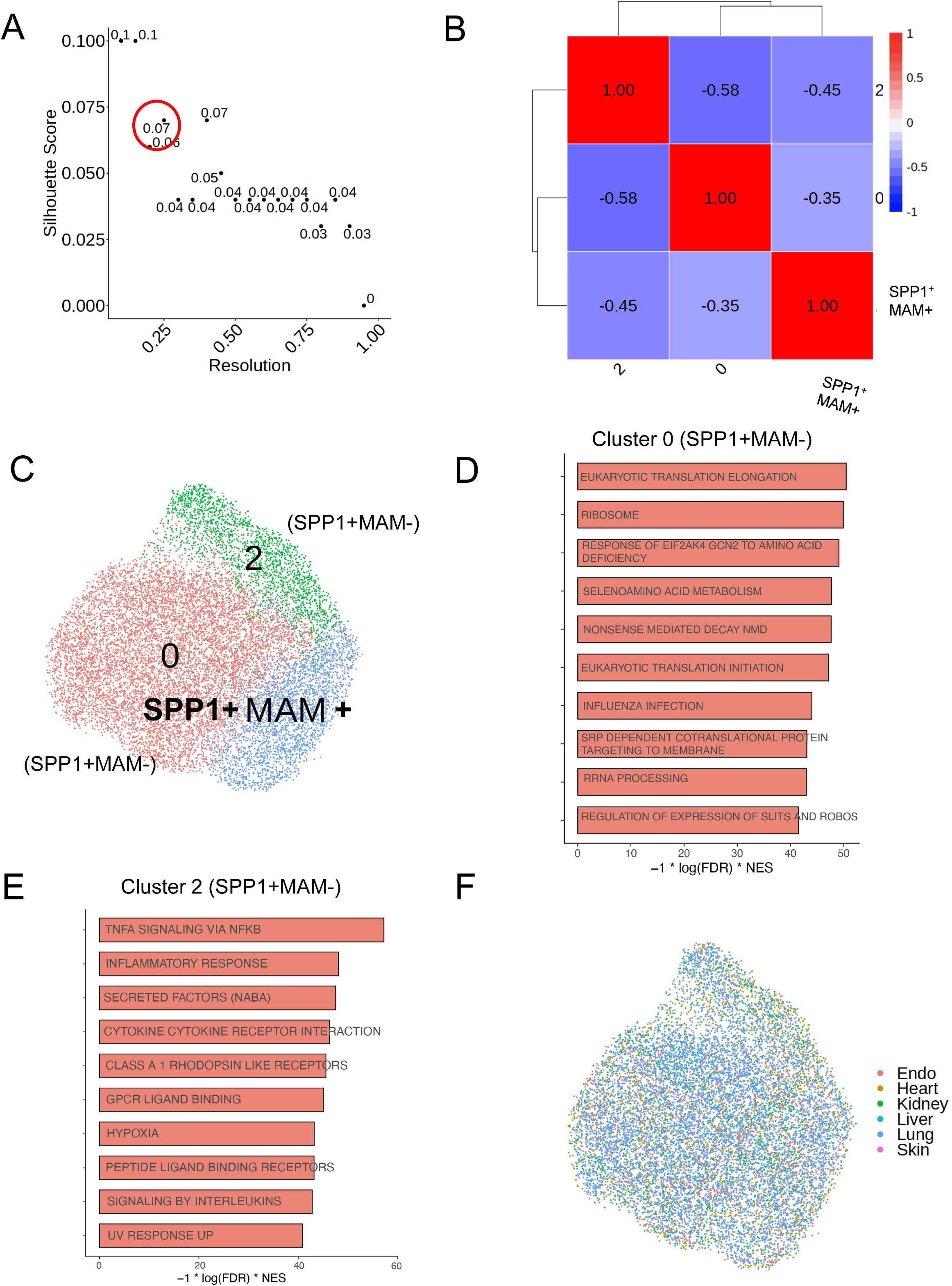
Data-driven clustering of SPP1**+** macrophages identifies a matrisome-associated macrophage polarization state. (**A**) Scatterplot of Silhouette score of single-cell clustering results by Seurat’s Louvain clustering algorithm at different resolutions (“res”) (see Methods). A res > 0.2 maximizes the Silhouette score (circled in red); res = 0.25 corresponds to Silhouette score of 0.07. (**B**) Heatmap showing the Spearman correlation between the pseudo-bulk profiles of clusters derived by using the selected res. Each box is labeled by the Spearman correlation between the cluster pair. (**C**) UMAP projection of SPP1+ macrophages pooled across all tissues (see also Table 1), with each cell colored by its cluster assignment. (D-E) Barplot summarizes GSEA results of differentially expressed genes between cluster 2 (D) and cluster 0 (**E**) (SPP1+MAM-) with respect to the remaining SPP1+ macrophages. The top 10 enriched annotation terms are shown alongside the strength of enrichment for each pathway. Functional enrichments for the SPP1+MAM+ cluster are reported in Figure 2D-E, and in Supplementary Table 5. (**F**) UMAP projection of SPP1+ macrophages, colored by tissue origin of each cell (see Table 1 for all the human organs/tissues used for the meta-analysis of SPP1+ macrophages).

**Supplementary Figure 8.**
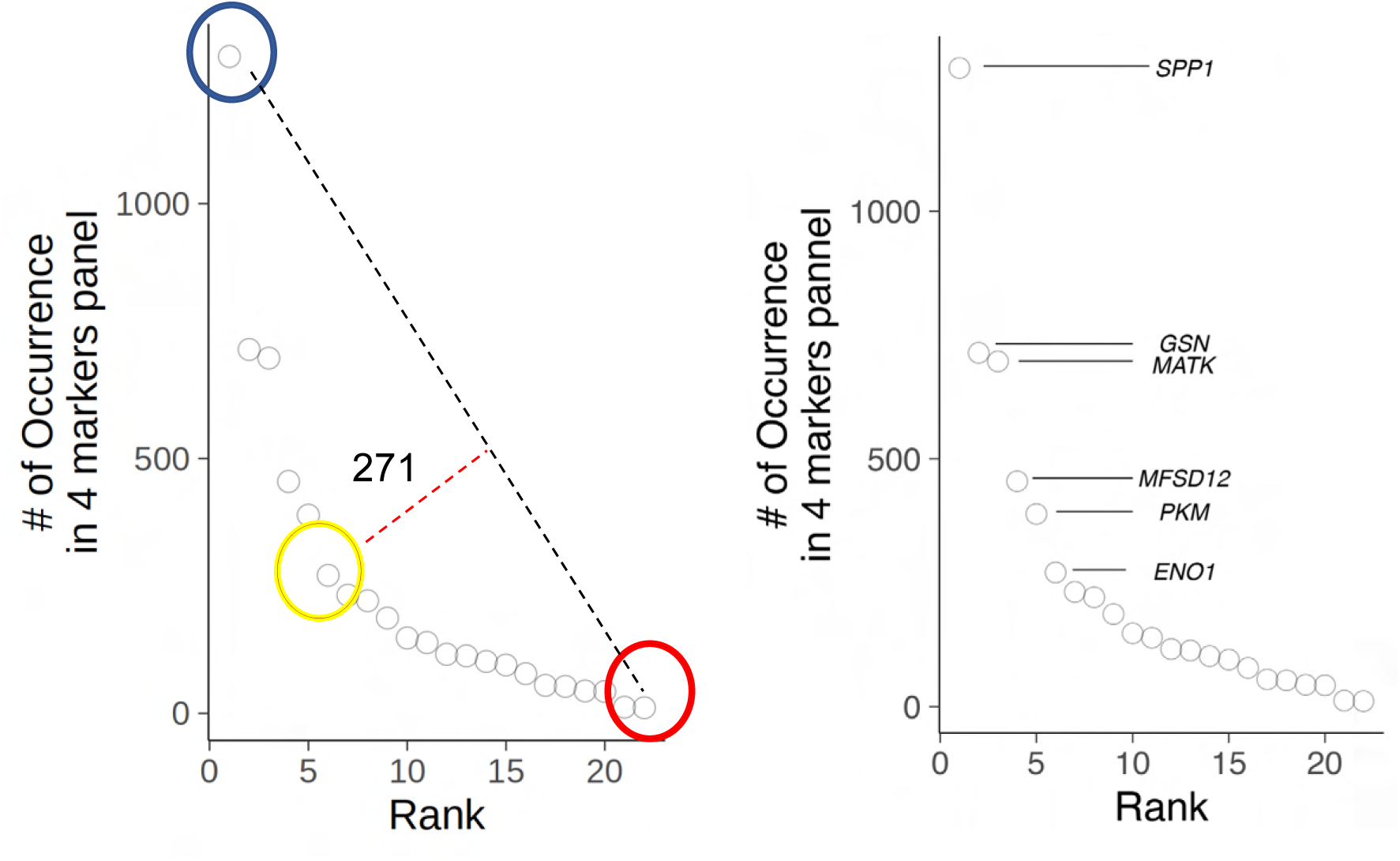
Markers of SPP1+MAM+ as predicted by COMET. (*left panel*) Scatterplot ranks the number of times a gene occurs in a 4 markers panel of SPP1+MAM+ macrophages as predicted by COMET (see Methods). The Elbow method was employed to select the top markers for SPP1+MAM+. The number of genes to be retained (starting from the most frequently occurring gene SPP1, see *right panel*) is chosen at the elbow, where a drop in the “number of occurrences in 4 markers panel” occurs on the scatterplot. To identify the elbow, a line connecting the first (blue circled) and the last (red circled) point on the scatter plot was drawn, and the orthogonal distance of each point to the line was calculated. The elbow of the plot is determined to be the point with the largest orthogonal distance to the line. The gene represented by this point on the scatterplot and other genes that occur more often than this gene in 4 markers panel predicted by COMET are retained as the marker genes of SPP1+MAM+ macrophages (*right panel*).

**Supplementary Figure 9.**
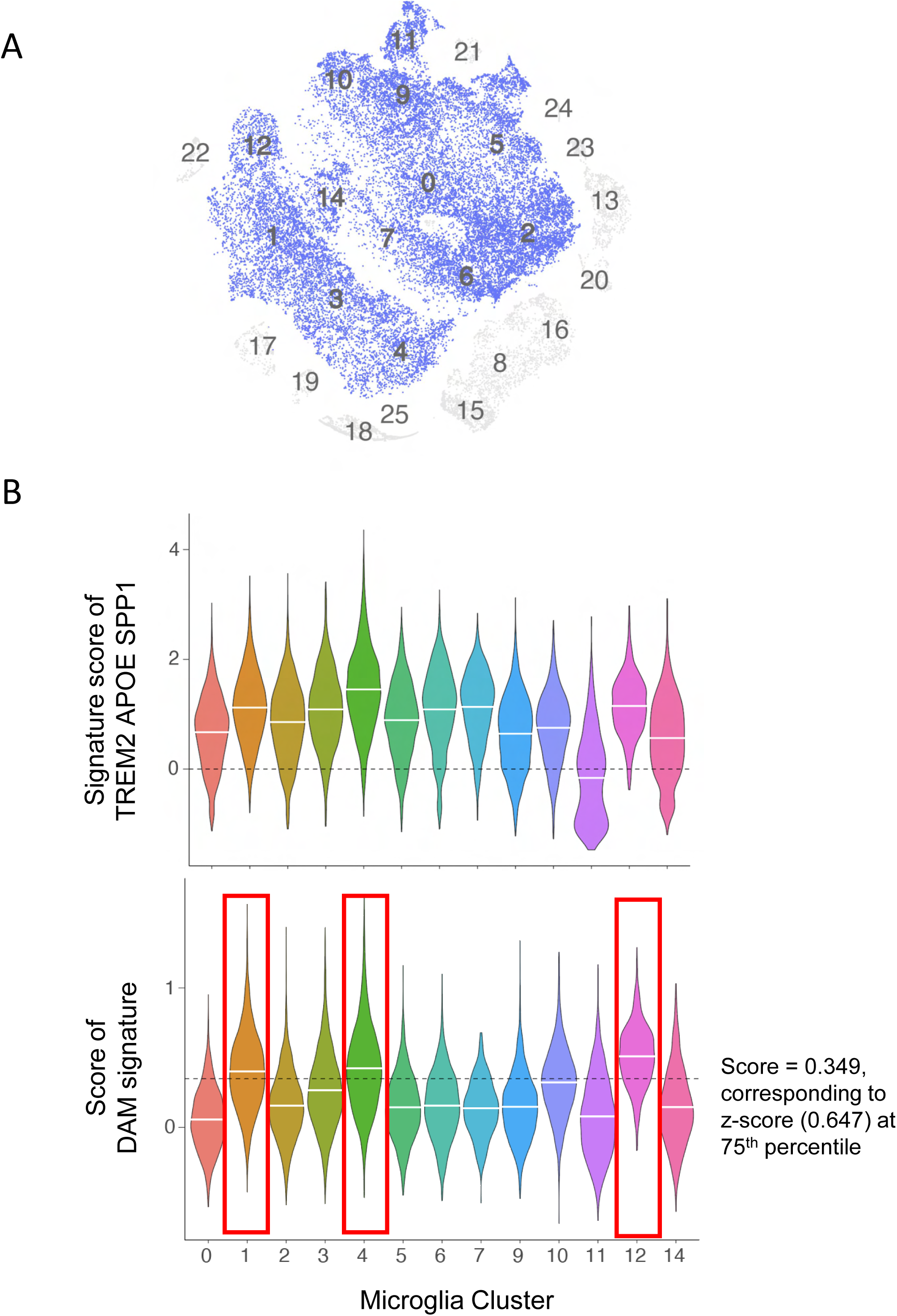
Derivation of disease-associated microglia (DAM) transcriptomic signature from the temporal lobe of epilepsy patients. (**A**) UMAP of immune cells isolated from the temporal lobe epilepsy patients (Kumar, et al., 2022 *Nature Neuroscience*; [Ref 119] in the manuscript; n = 4 patients). Microglia as defined in the original paper are colored in blue; n = 5,645 cells. (**B**) Violinplot displays the expression of *TREM2, APOE*, and *SPP1* (*top*) and the DAM signature within the microglia (*bottom*). Median of the expression/score in each microglia cluster is indicated with a white line on the violin. Dashed line marks the DAM signature score at the 75th percentile by z-score (*bottom*). Clusters 1, 4 and 12 are classified as “DAM clusters”.

**Supplementary Figure 10.**
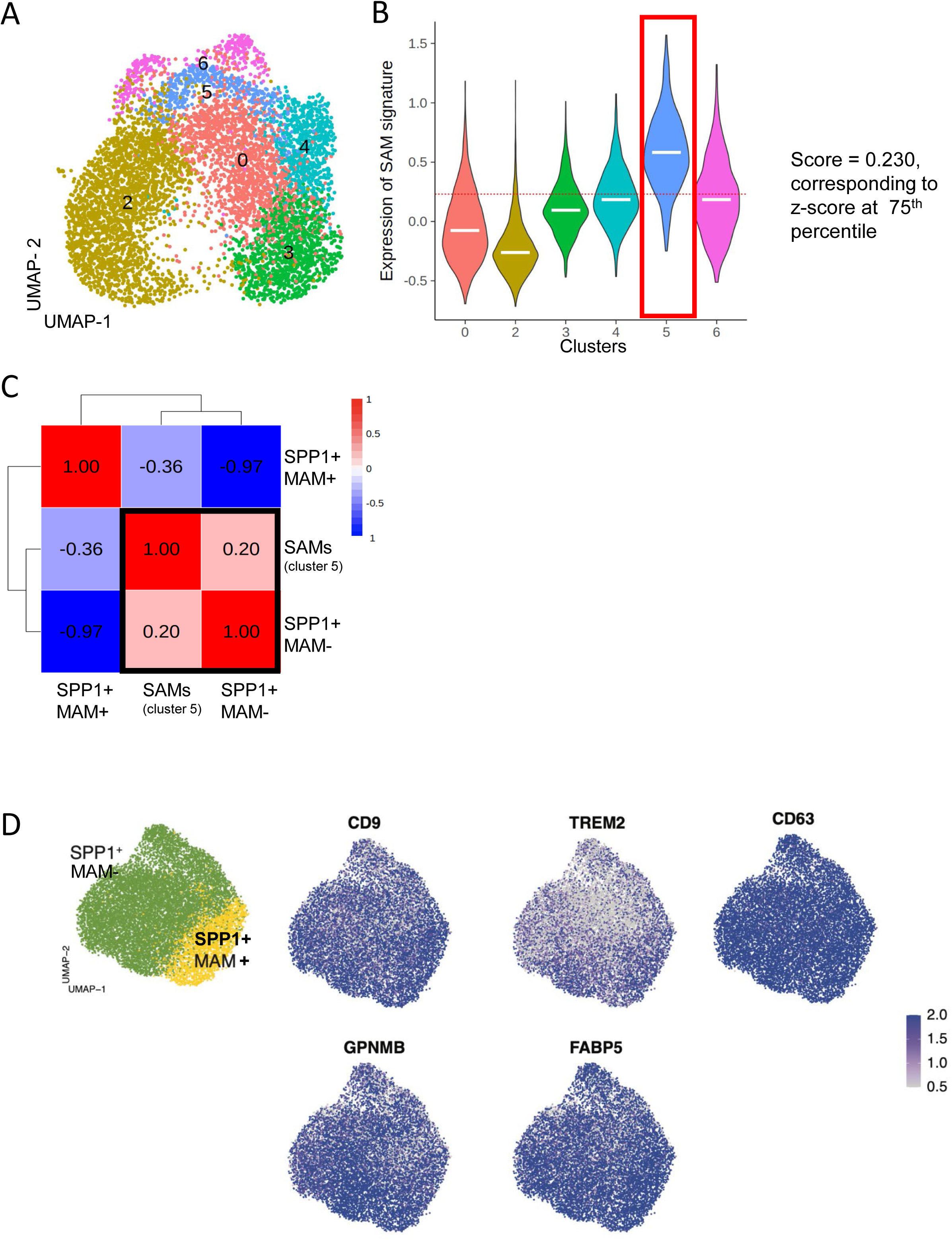
Scar-associated macrophages (SAMs) show weak transcriptomic similarity with SPP1+MAM- macrophages. (**A**) UMAP of monocytes/macrophages derived from human cirrhotic and control livers (see Supplementary Figure 2 for data-driven derivation of clusters). (**B**) Violin plots showing transcriptomic signature of SAMs (as defined by Ramachandran et al., [Ref 25]) for each cluster defined in Supplementary Figure 2. See Supplementary Table 2 for genes that belong to this signature. Median of the expression/score in each cluster is indicated with a white line on the violin. Dashed line marks the SAM signature score at the 75th percentile by z-score. Cluster 5 is classified as “SAM clusters”. (**C**) Heatmap displays the Spearman correlation between pseudo-bulk profiles of SPP1+MAM+, SAMs (as defined by Ramachandran et al.; [Ref 25]) and SPP1+MAM- macrophages. The pseudo-bulk profiles were constructed using the top 1,000 highly variable genes common to the above-mentioned macrophage clusters. (**D**) UMAP of SPP1+MAM+ and SPP1+MAM- macrophages, where each cell is colored by expression of marker genes of SAMs defined by Fabre, et al. (Fabre, et al, 2022 *Biorxiv;* [Ref 34]).

**Supplementary Figure 11.**
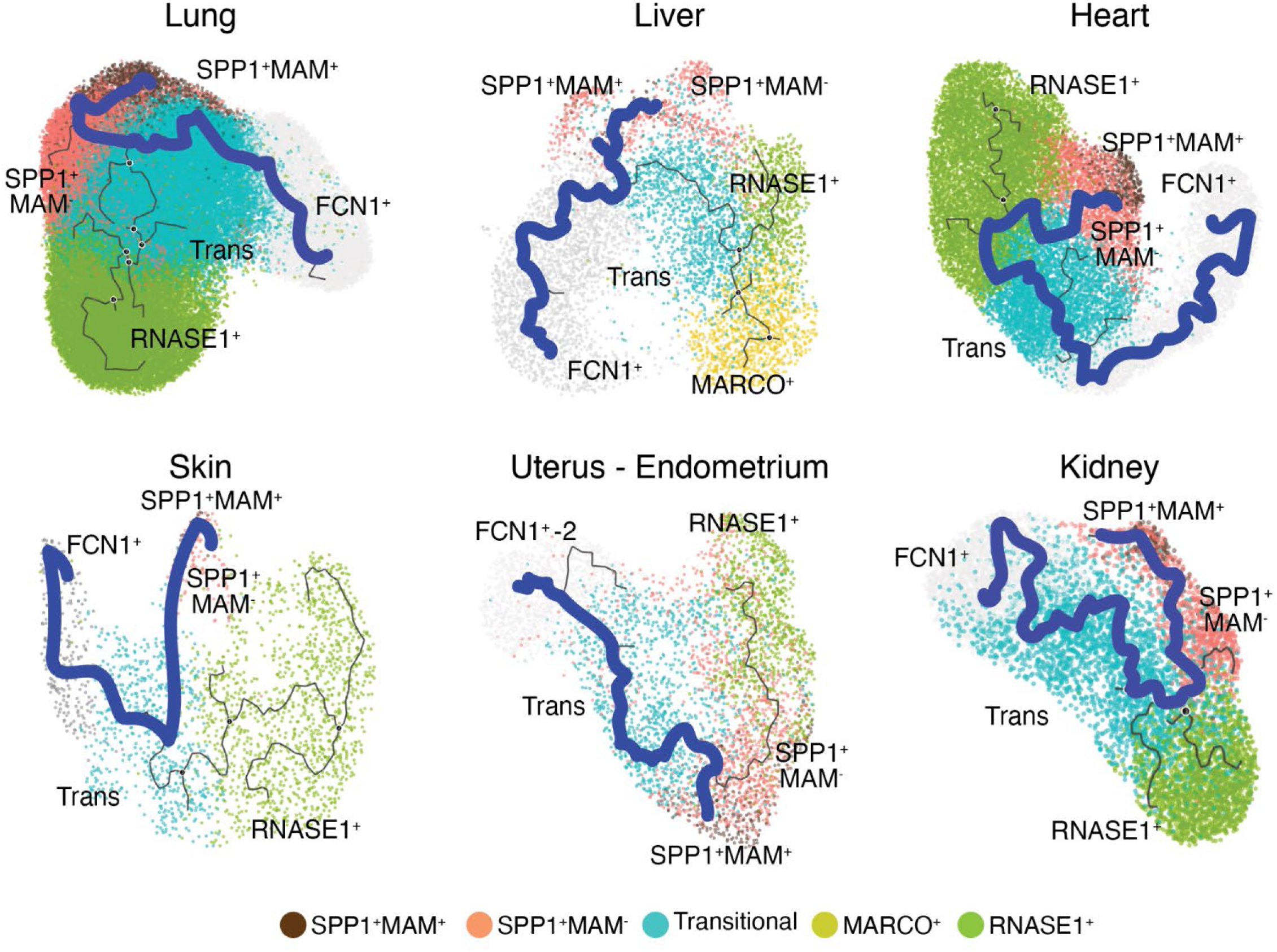
Monocle analysis recapitulates the conserved differentiation trajectory from FCN1^+^ monocytes to SPP1+ macrophages across multiple tissues. Monocle differentiation trajectory analyses in the lung, liver, heart, skin, endometrium, and kidney (see Table 1 for information regarding the human tissues). The predicted trajectories by Monocle are based on UMAP projection of macrophages in each organ (black line). The differentiation trajectories from FCN1^+^ monocytes to SPP1+ macrophages are shown with a thick blue line.

**Supplementary Figure 12.**
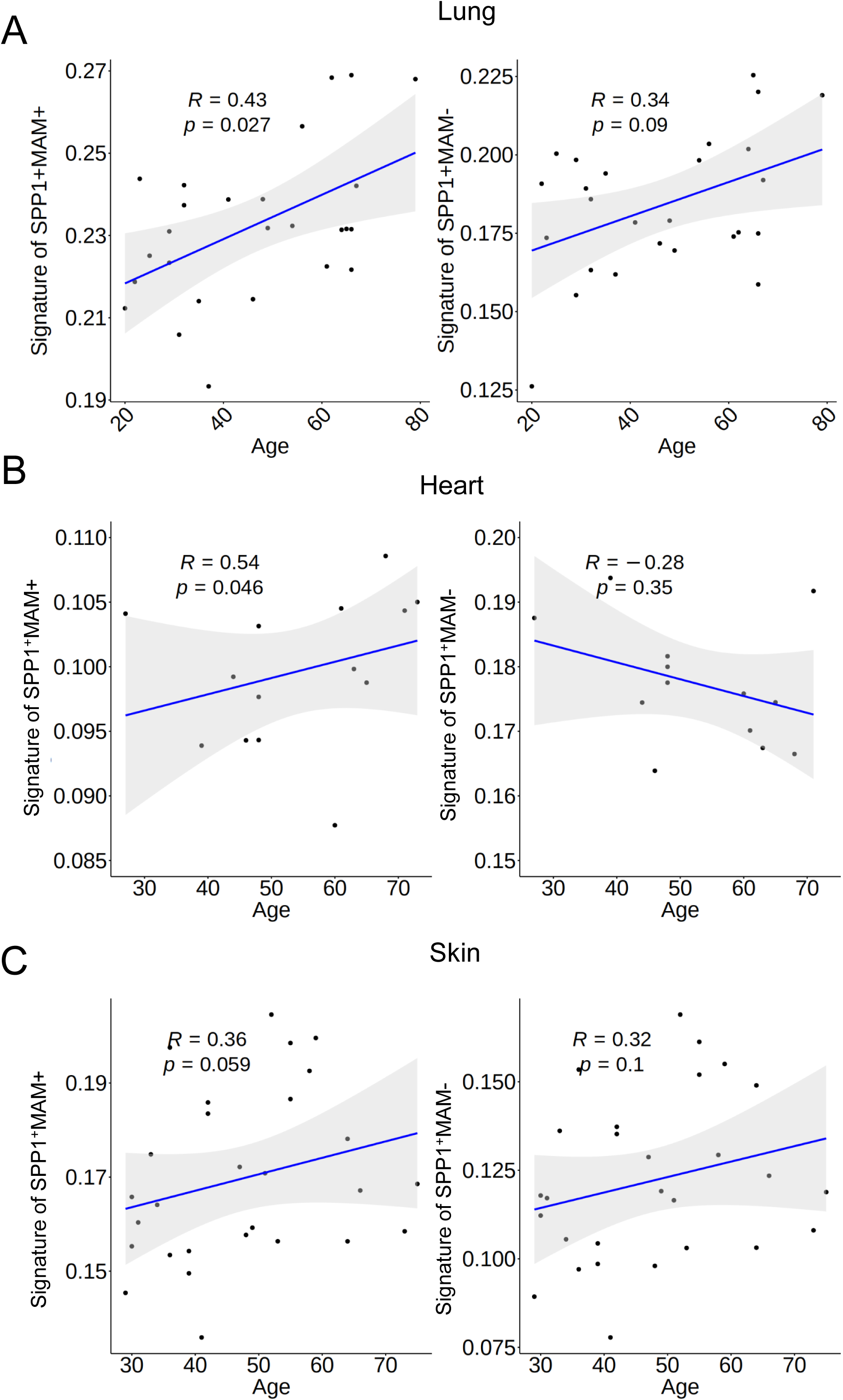
Transcriptomic signature of SPP1+MAM+ is increased with age in human macrophages from lungs, heart, and skin during homeostasis. (**A-C**) Scatter-plot displays the pseudo-bulk expression of SPP1+MAM+ signature (*left*) and SPP1+MAM- signature (*right*) in macrophages from human lungs (**A**), heart (**B**) and skin (**C**) against age, where each dot represents a healthy human sample (lung, n = 26; heart, n = 14; skin, n = 28). A linear regression was used to model the relationship between SPP1+MAM+ or SPP1+MAM+ signatures and age (plotted in blue line, with gray band indicating 95% confidence interval of the regression line). The non-parametric Spearman correlation *R* (and associated significance P-value) are also indicated within in each plot.

**Supplementary Figure 13.**
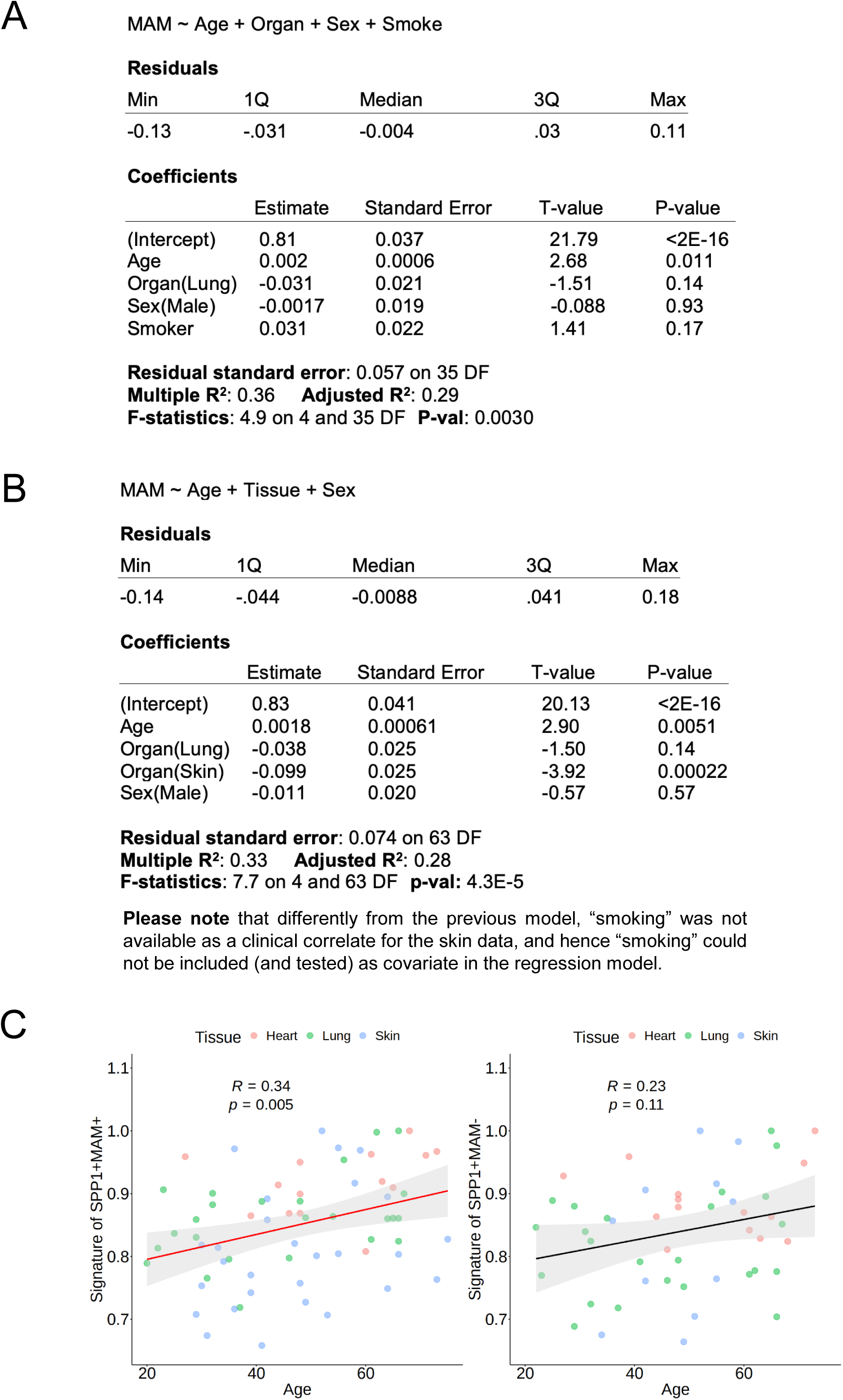
SPP1+MAM+ are associated with aging. (**A-B**) Linear regression analysis of SPP1+MAM+ transcriptomic signature against tissue, gender and smoking status using pseudo-bulk expression profiles of human tissue SPP1+MAM+ macrophages (the dependent variable (*Y*) in the regression model) from healthy heart and lungs (**A**) and from heart, lung and skin (**B**). (**C**) Scatterplot showing the change in transcriptomic signature of SPP1+MAM+ (*left*) or SPP1+MAM- (*right*) against age, with each dot representing a human sample. Spearman correlation between pseudo- bulk expression profiles of the transcriptomic signatures and age in lung (n = 26), heart (n = 14) and skin (n = 28). The correlation coefficient and its statistical significance (P-value) are shown within each plot.

**Supplementary Figure 14.**
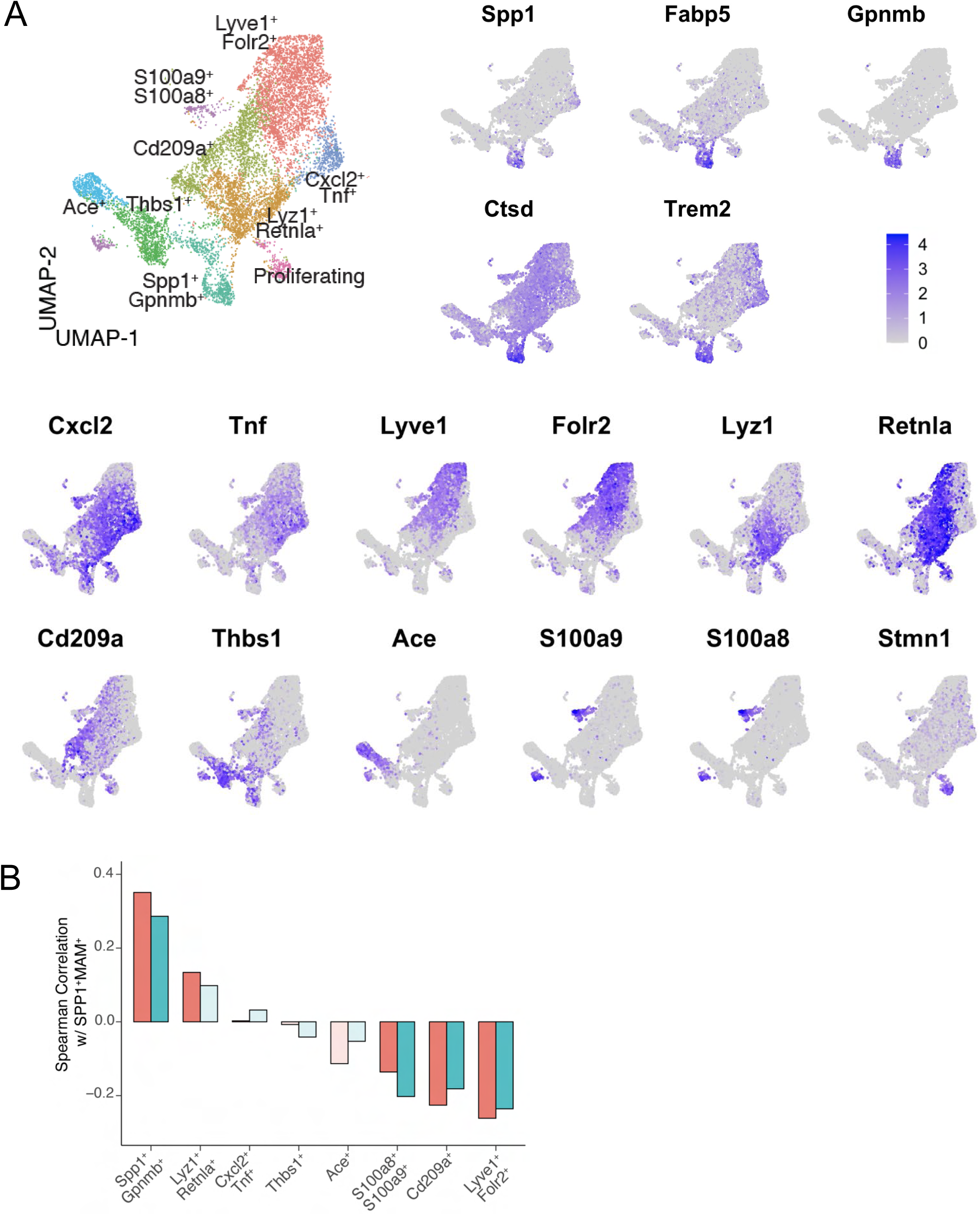
Deriving aging-related *Spp1*+*Gpnmb*+ Cl6 cluster from scRNA-seq of aging macrophages from murine skeletal muscle. (**A**) UMAP representation of sorted skeletal macrophages (*Cd11b*+, *F4/80*+) from old (23 months, n = 3) and young (3 months, n = 3) mice (Krasniewski, et al, 2022 *Elife*) (young, n = 4,200 cells; old, n = 5,222 cells), with each cell colored by the marker genes of macrophage cluster in the dataset as indicated by Krasniewski et al. The clusters were derived using the R code provided by the authors in the original publication. (**B**) Barplot displays the Spearman correlation (*y-axis*) between the pseudo-bulk profile of each skeletal macrophage cluster defined by Krasniewski et al (*x-axis*) and the human SPP1+MAM+ macrophages in this dataset, stratified by age group. Young: turquoise, old: salmon. Murine skeletal macrophage clusters with significant correlations (P-value<0.05) with SPP1+MAM+ macrophage signature are indicated by filled color bars.

